# Comparative transcriptome analysis of tomato cultivars pretreated with Laminarin reveals differential expression of defense-responsive genes against Early blight disease

**DOI:** 10.1101/2025.10.03.680217

**Authors:** Govindan Muthukumar, Sowmiyarithika Sowrirajan Padmavathy, Jeyapandi Mohana Prasad, Chinnasamy Arulvasu, Nallamuthu Godhantaraman, Mehanathan Muthamilarasan, Nagarathnam Radhakrishnan

## Abstract

Eco-friendly measures to control disease progression are the need of the hour since chemical fungicides have been harmful to the environment. This study observed that the algal polysaccharide Laminarin triggers a defense response against early blight disease in both tolerant and susceptible tomato cultivars. A dose-dependent reduction in cell death was observed in tomato leaves pre-treated with 0.1% Laminarin and then infected with Alternaria solani in a susceptible and tolerant cultivar. As a result, 0.1% Laminarin was chosen for transcriptome analysis. Transcriptome analysis revealed the upregulation of defense-related genes, such as EF-hand domain-containing protein, Heat shock protein 70, Heat shock factor (HSF) type DNA binding domain-containing protein, Non-specific serine/threonine protein kinase, AP2/ERF domain-containing protein, Leucine-rich repeat-containing N-terminal plant type domain-containing protein, and Subtilisin-like protease. The study also validated the differentially expressed genes using qRT-PCR for bZIP, nsLTP, SKP1, and V-type ATPase genes. Overall, the study demonstrated that the algal polysaccharide Laminarin could induce defense-related genes and provide resistance against early blight disease in tomato plants.

**KEY MESSAGE:** Tomatoes defense-responsive genes showed differential expression in tolerant and susceptible cultivars when treated with laminarin, followed by *Alternaria solani* infection.

## INTRODUCTION

Non-motile plants continuously encounter many harsh environmental conditions during their growth phase. *Solanum lycopersicum* Linn. (Tomato) is one of the world’s most desirable economic and extensively marketing vegetable crops and belongs to the family of Solanaceae (Khan *et al*., 2014). Tomato plants offer a straightforward gene pool that is enough for examining the genes involved in the resistance to biotic and abiotic stress (Bai *et al*., 2018). Due to its short lifespan and several benefits for genomic, transcriptomic, metabolomic, and phenotypic studies, the tomato is one of the most thoroughly studied agricultural plants (Ranjan *et al*., 2012; Lin *et al*., 2014). Analysis of plant-fungal interactions allows the development of various new strategies for enhancing the production and protection of various crops in agriculture (Zeilinger *et al*., 2016). Among pathogens, fungi inflict the most significant losses, as high as 42%, followed by bacteria (27%), viruses (18%), and nematodes (13%) (Khan & Jairajpuri, 2010).

Induced resistance in plants, using natural compounds is a significant and eco-friendly alternative approach to agrochemicals in controlling plant diseases such as bacteria, viruses, and fungi. Induced resistance can boost the host-natural defense responses against different diseases (Van Loon *et al*., 2000; Boubakri, 2020). Two types of induced resistance involved in plants, known as systemic acquired resistance (SAR) and induced systemic resistance (ISR), occur when a plant’s defenses are preconditioned by an earlier infection or any treatment that results in resistance against subsequent challenge by a pathogen (Choudhary *et al*., 2007). SAR is a form of induced resistance by prior treatment with elicitors from non-pathogenic, avirulent or virulent microbes, as well as elicitor molecules derived from natural sources and synthetic chemical stimuli, such as sodium alginate (Dey *et al*., 2019), kappa carrageenan (Mani *et al*., 2021), salicylic acid (Radhakrishnan & Balasubramanian, 2009), sulfamethoxazole (Thinakaran *et al*., 2020) and methyl jasmonate methyl ester (Kamakshi *et al*., 2023). Induced systemic resistance (ISR) is triggered by pathogen infection. Its mode of action does not depend on direct killing or inhibition of the pathogen infection but rather on strengthening the chemical or physical barriers of the host plant (Ryals *et al*., 1996). Plant pathogen interactions plants trigger a salicylic acid-dependent signaling cascade, leading to systemic resistance against various pathogens (fungi, bacteria, and viruses) (Kamle *et al*., 2020).

Early blight disease of tomatoes caused by *Alternaria solani* is the most prevalent and destructive disease of tomatoes. It significantly reduces the amount and quality of fruit supply (Rashmi *et al*., 2012). Early blight disease is a foliar infection; thus, it can cause total defoliation in severe situations. Early blight affects every part of the tomato plant above ground, and depending on the symptoms, it can be divided into three different phases or stages: collar rot in seedlings (cankers), early leaf blight, and sunken lesions in fruit (rot) (Sherf *et al*., 1986; Foolad *et al*., 2002). The tomato plant’s leaf blight stage or phase can be seen at earlier stages, but it usually appears during the adult stage. Leaf blight appears as small, dark, and coalescing concentric ring-like lesions, often called “bullseye” patterned leaf spots (Adhikari *et* al., 2017; Rotem, 1994). The leaf tissue surrounded by yellow lesions leads to the death of the leaf. This pathogen causes disease by producing toxic secondary metabolites such as alternaric acid and zinniol, which are not host-specific. These toxins are released into the lesion site of leaves, causing damage to nearby cells, leading them to die off or senesce (break down) (Lawrence *et al*., 2000). As the disease progresses, it causes leaves to dry, leading to loss of foliage (Adhikari *et* al., 2017). Currently, early blight disease is controlled mainly by the application of agrochemicals or chemical fungicides. An effective control measure for early blight disease of tomatoes involved spraying mancozeb (86.4%), effectively reducing the suppressing symptoms in the Pusa Ruby tomato cultivar.(Maheswari *et al*., 1991; Gondal *et al*., 2012; Kumar *et al*., 2013; Sadana & Didwania, 2015). Other fungicides documented to reduce disease symptoms include carbendazim (76.67%), antracol (62.69%), kavach (50.74%), insignia (54.63%), copper oxychloride, pyroclostrobin and hexaconazole effective inhibition of mycelial growth (Kumar *et al*., 2017).

Elicitors are substances that cause signals that plants detect and either trigger or induce defensive responses (Wiesel *et al*., 2014). Although few elicitors used to induce plant defense are a natural source, many synthetic products such as jasmonic acid (JA), salicylic acid (SA), beta-amino butyric acid (BABA), and benzothiadiazole (BTH) are also widely in practice (Radhakrishnan & Balasubramanian, 2009; Wiesel *et al*., 2014; Bektas & Eulgem, 2015). These agrochemicals activate plants crop’s defense against many pathogens (Bektas & Eulgem, 2015; Burketova *et al*., 2015; Poveda, 2020). Besides, elicitors of biotic origin can induce structural or biochemical responses associated with the expression of plant disease resistance. Brown alga is one of the rich sources of biopolymer laminarin, a water-soluble small storage polysaccharide previously reported to be an efficient elicitor against pathogens (Shukla *et al*., 2021). Laminarin has been shown to enhance resistance against *Fusicladium oleagineum* infection in olive trees. (Tziros *et al*., 2021). In *Camellia sinensis* (Tea plant), Laminarin regulates SA, ABA, MAPK signaling, and H_2_O_2_ against tea leafhoppers (Xin *et al*., 2019). In *Arabidopsis*, Laminarin was shown to induce plant defensin-like 202 (DEFL202), stabilizing the chloroplast under salt stress and enhancing plant growth activity under heat stress (Wu *et al*., 2016). Laminarin has demonstrated its ability to boost resistance in *Vitis vinifera* (grapevine) and trigger ROS-dependent defense mechanisms against *Plasmopara viticola*.(Gauthier *et al*., 2014).

To understand the molecular networks involved in plant/host-pathogen interactions, we can analyze gene expression through transcriptomic analysis. This can help identify essential genes for pathogen resistance. (Vega *et al*., 2015; Campos *et al*., 2021). The present study’s transcriptome analysis of tomato pretreated with elicitor laminarin and *Alternaria solani* infection has shed light on the cross-talking among different signaling pathways.

## MATERIALS AND METHODS

### Plant material, Elicitor, and Pathogen Culture

The seeds of a susceptible tomato cultivar, PKM1, were procured from Horticultural College and Research Institute, Periyakulam, Tamil Nadu, India, and an early blight disease-tolerant tomato cultivar, Arka Rakshak, was procured from the Indian Institute of Horticultural Research, Bengaluru, Karnataka, India. Thirty-day-old healthy plants were foliar sprayed with the elicitor, Laminarin (LaM) (YL76431-Biosynth ® Carbosynth, United Kingdom) at a concentration of 0.1% (w/v) on adaxial and abaxial leaves. The control plants were sprayed with sterile, double-distilled water. Both the cultivars were inoculated with the fungal pathogen *Alternaria solani* (2 x 10^5^ conidia per mL) after 48 hours of elicitor treatment. All the plants were kept in controlled conditions at 25 °C, with a photoperiod of 16 h light/8 h dark, until the first visible symptoms post-infection appeared. Leaf samples of susceptible and resistant tomato cultivars were harvested 48 hrs after the first visible symptoms of *A.solani* infection. Control and treated plants were collected for each variety to compare gene expression.

### Cell death analysis

Cell death analysis of experimental leaves was monitored by following the method of Levine et al. (1994). A known quantity of leaves showing characteristic browning symptoms were ground with one ml of sterile distilled water. For each sample, a 500μL aliquot of sample was incubated with 0.05% Evans blue for 30 minutes and then washed extensively. The dye bound to dead cells was solubilized in 50% (v/v) methanol with 1% (w/v) Sodium dodecyl sulfate (SDS) for 30 min at 50°C and quantified by A600.

### RNA extraction and Illumina sequencing

Total RNA was extracted from all the experimental leaves with TRI reagent (Sigma-Aldrich, USA) by following the procedure described by Mani *et al*. (2021). The quality of RNA was ascertained by resolving on a denaturing 1.2% (w/v) formaldehyde gel. Nucleic acid quality and quantity were determined with a NanoDrop 1000 Spectrophotometer (Thermo Scientific, USA). Samples with an A260/A280 ratio of 1.8-2.0 and RIN values above 7.0 were used for sequencing. Sequencing was performed using an Illumina HiSeq 2500 platform. The resultant fastq files were checked for base and sequence quality score distributions, average base content per read, GC distribution in the reads, PCR amplification issues, and over-represented sequences (Mani *et al*., 2021).

### Pre-processing, alignment, and differential expression analyses

The raw reads were processed using the AdapterRemoval (version 2.2.0) tool to trim adaptor sequences and remove low-quality bases. The ribosomal RNA sequences were removed by aligning the reads with the SILVA database using bowtie2 (version 2.2.9) and subsequent workflow using SAMtools (version 1.3.1), Sambamba (version 0.6.7) and BamUtil (version 1.0.13). The reads were then aligned to the tomato genome, downloaded from Ensembl Plants (ftp://ftp.ensemblgenomes.org). The alignment was performed using the STAR program (version 2.5.3a). After aligning the reads with transcriptome, differential expression analysis was performed using the Cuffdiff program of the Cufflinks package (version 2.2.1) at p-value cut off 0.05 and 0.01 along with log2 fold change cut-off 2 (Mani et al., 2021).

### Validation of quantitative RT PCR analysis

The total RNA was isolated, and the first-strand cDNA synthesis was performed using the PrimeScript 1st-Strand cDNA Synthesis Kit (Takara) following the manufacturer’s instructions. Gene-specific primers for qRT-PCR analysis were designed using the NCBI Primer-BLAST with default parameters. The qRT-PCR analysis was performed using SYBR Green detection chemistry on a 7900HT Sequence Detection System with software version 2.3 (Applied Biosystems). Reactions were carried out in a final volume of 20 µl. The components included 2 µl of 5X diluted cDNA, 250 nM of each primer, and 10 µl of TB Green Premix Ex Taq II (Takara). The reaction mixture was initially denatured for 2 min at 50 °C, followed by 10 min at 95 °C, 40 cycles of 15 s at 95 °C, and 1 min at 60 °C. Furthermore, to ensure the amplification specificity and absence of multiple amplicons or primer dimers, melting curve analysis (60 to 95 °C after 40 cycles) and agarose gel electrophoresis were performed. Three technical replicates for three independent biological replicates were maintained, and the transcript abundance was normalized to the internal control Elongation Factor 1α. The analysis was conducted using the 2^-ΔΔCt^ method. (Mani *et al*., 2021).

### Statistical analysis

All experimental data were obtained from three replicates and the data were expressed as the mean ± standard error of the mean. Statistical analysis was performed using Student’s t-test to determine the significant differences at p < 0.01 and p < 0.05. All statistical analyses were performed using the OriginPro software (OriginLab Corporation, Northampton,USA).

## RESULTS AND DISCUSSION

During plant-pathogen interaction, pathogens produce various chemicals to suppress the defense system of the host plant. The recognition of an invading pathogen triggers a signal transduction pathway that results in the coordinated activation of many plant defense mechanisms, including the accumulation of salicylic acid, the synthesis of pathogenesis-related proteins, the thickening of cell walls and the increased production of antimicrobial compounds (phytoalexins) as well as ROS leading to cell death (Durrant & Dong, 2004). In plants, two types of induced resistance activate the defense mechanism: Systemic Acquired Resistance (SAR) and Induced Systemic Resistance (ISR). The SAR pathway is activated due to prior pathogen infection, leading to the involvement of the salicylic acid-dependent SAR pathway in combating necrotrophic pathogens (Panpatte *et al*., 2020). Several algal polysaccharides have been utilized as elicitors against various fungal diseases in plants and have been demonstrated to trigger plant resistance against numerous pathogens, including fungi (Dey *et al*., 2019; Mani *et al*., 2021; Shukla *et al*., 2021). Various chemicals, insecticides, and fungicides have been used to remove fungal infestations. Therefore, the alternative strategy is to use biopolymers to induce plant defense as they are eco-friendly and economical for sustainable agriculture. Among them, Laminarin has proven to reduce the disease severity very effectively and induce plant resistance in grapevine against *Plasmopara viticola* (downy mildew) (Adrian *et al*., 2017), grapevine powdery mildew (Pugliese *et al*., 2018), olive leaf spot disease caused by *Fusicladium oleagineum* in olive tree (Tziros *et al*., 2021), wheat against *Zymoseptoria tritici* (de-Borba *et al*., 2022) and wild tomato species *S. chilense* against *Phytophthora infestans (*Kahlon *et al*., 2023). Different groups of seaweed are used as natural biostimulants or elicitors to activate various immune pathways and the activity of defense enzymes (Dey *et al*., 2019; Mani *et al*., 2021; Shukla *et al*., 2021).

### Cell death analysis

Cell death analysis of all the experimental leaves was studied using Evans blue dye. In the susceptible tomato cultivar infected with *A. solani*, maximum cell death was recorded when compared to the tolerant tomato cultivar and control groups. There was a time-dependent increase in cell death recorded in the susceptible cultivar from 0 to 6 hrs in the tolerant cultivar. In comparison, there was slow progression in cell death from 0 to 12 hrs. These results confirmed that *A. solani* infection resulted in differential cell death in tomato leaves. In Laminarin (0.010, 0.025, 0.050, 0.075, and 0.10%) pretreatment followed by *A. solani* infection resulted in a reduction in cell death due to reduced colonization of the pathogen. Laminarin (0.010%) was pretreated and followed by *A. solani* infection of tomato leaves of both cultivars, which showed higher cell death compared to other concentrations tested (**Fig. 1**.). Compared with control groups, Laminarin (0.1%) effectively reduced cell death in susceptible cultivars. Laminarin (0.1%) pretreated reduces cell death in susceptible and tolerant cultivars. A dose-dependent reduction in cell death was observed in tomato leaves pre-treated with 0.1% Laminarin and then infected with *A. solani* in a tolerant cultivar and as a result, 0.1% Laminarin was chosen for transcriptome analysis.

**Fig. 1.**
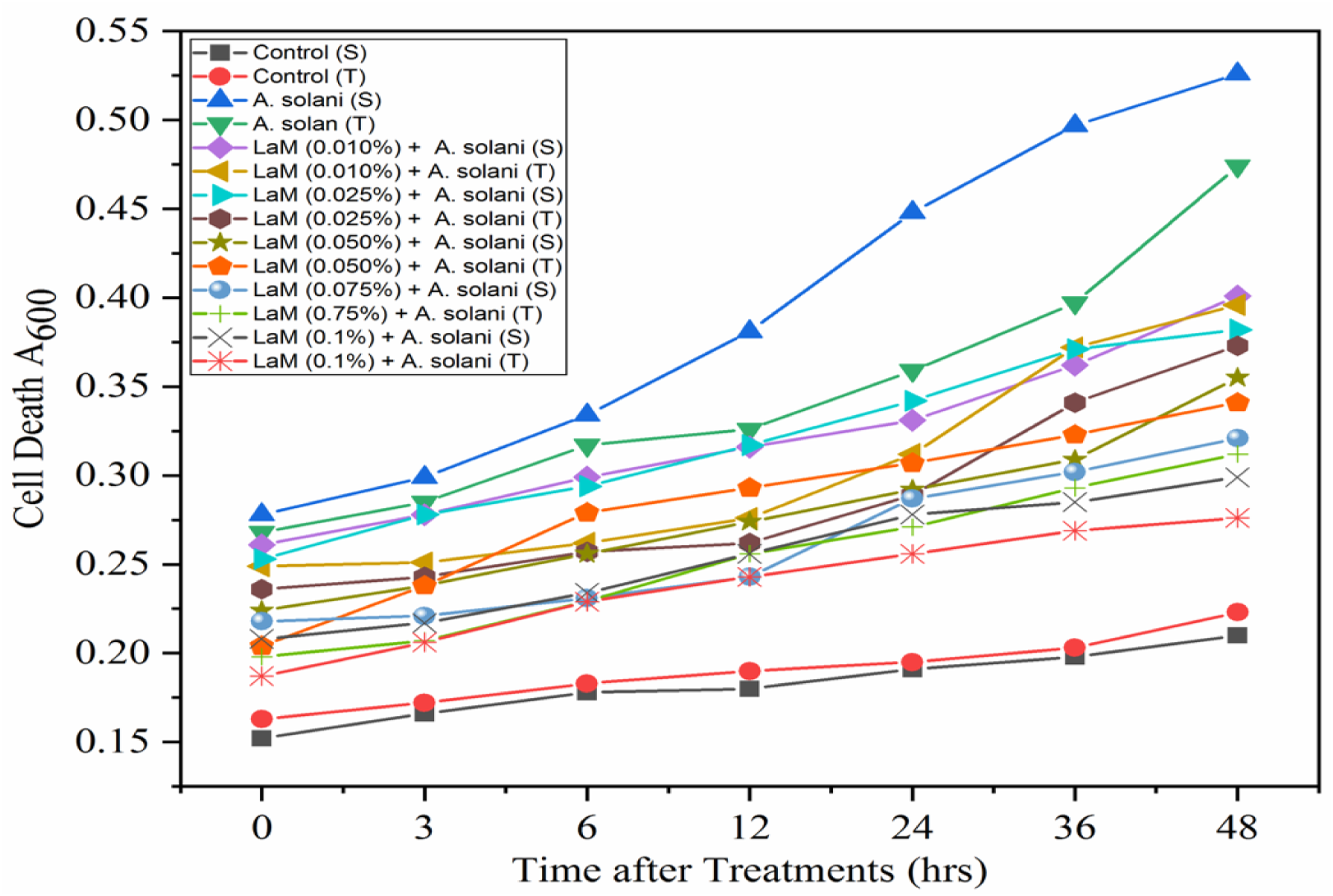
Cell Death Analysis of leaves of *S. lycopersicum* pretreated with Laminarin and infected with *A. solani.*1. Control (S), 2. Control (T), 3. *A. solani* infected (S), 4. *A. solani* infected (T), 5. Laminarin (0.010%) pretreated and followed by *solani* infected (S), 6. Laminarin (0.010%) pretreated and followed by *A. solani* infected (T), 7. Laminarin (0.025%) pretreated and followed by *A. solani* infected (S), 8. Laminarin (0.025%) pretreated and followed by *A. solani* infected (T), 9. Laminarin (0.050%) pretreated and followed by *A. solani* infected (S), 10. Laminarin (0.050%) pretreated and followed by *A. solani* infected (T), 11. Laminarin (0.075%) pretreated and followed by *A. solani* infected (S), 12. Laminarin (0.075%) pretreated and followed by *A. solani* infected (T), 13. Laminarin (0.1%) pretreated and followed by *A. solani* infected (S), 14. Laminarin (0.1%) was pretreated and followed by *A. solani* infected (T), S - Susceptible cultivar and T - Tolerant cultivar.

### Transcriptome Analysis

In the present study, RNA sequencing was conducted to identify *Solanum lycopersicum* genes involved in the tomato - *Alternaria solani* interaction and to study the elicitor laminarin-induced genes in tomato*es.* The RNA sequencing data was used to identify differentially expressed genes responding to pathogen infection and laminarin treatment in tolerant and susceptible tomato cultivars. A total of eight samples such as Control (Susceptible - S), control (Tolerant - T), *A. solani* (S), *A. solani* (T), Laminarin (0.1%) (S), Laminarin (0.1%) (T), Laminarin (0.1%) (S) + *A. solani* (S), Laminarin (0.1%) (T) + *A. solani* (T) were used to produce RNA-seq reads using Illumina RNA-seq deep sequencing. The RNA-seq generated transcriptome data from each library constructed from tolerant and susceptible cultivars. A total of 320761632 raw reads and 311463562 filtered reads (HQ) were generated, with 97.23% accuracy. After removing adaptors, low-quality reads, and contaminants, clean reads were obtained from each sample, of which 286753873 aligned reads with a percentage of 92.72 were found to be expressed in a total of all eight experimental samples (**Table 1**). The average aligned reads percentage in the libraries were 33097505 and 38590962 for tolerant and susceptible cultivars, respectively. Moreover, we conducted MA plots, volcano plots, Gene Ontology (GO), and Pathways enrichment analyses of the differentially expressed genes (DEGs) to reveal the transcriptomic characteristics of susceptible and resistant cultivars.

**Table 1.** Summary of filtered reads generated in RNA-Seq analysis of susceptible and resistant cultivars of tomato plants pretreated with laminarin and/or *A. solani* infection.

| S. No. | Sample name | Total Raw Reads | Total Filtered Reads (HQ) | HQ Reads Percentage | Aligned Reads | Aligned Percentage |
| --- | --- | --- | --- | --- | --- | --- |
| 1 | Control (S) | 53593904 | 52486310 | 97.93 | 47714536 | 90.91 |
| 2 | Control (T) | 31532704 | 30871098 | 97.90 | 29339109 | 95.04 |
| 3 | *A. solani* (S) | 28932592 | 27901736 | 96.43 | 25257792 | 90.52 |
| 4 | *A. solani* (T) | 54601440 | 51214640 | 93.79 | 41247172 | 80.54 |
| 5 | LaM (S) | 35785250 | 35185338 | 98.32 | 34038325 | 96.74 |
| 6 | LaM (T) | 32838670 | 32152264 | 97.90 | 31096314 | 96.72 |
| 7 | LaM + *A. solani* (S) | 50758160 | 49655086 | 97.82 | 47353198 | 95.36 |
| 8 | LaM + *A. solani* (T) | 32718912 | 31997090 | 97.79 | 30707427 | 95.97 |
| **Total & Average** |  | **320761632** | **311463562** | **97.23** | **286753873** | **92.72** |

### Transcriptional changes in tolerant and susceptible tomato cultivar upon *Alternaria solani* infection

#### Comparison of differentially expressed genes between control and *Alternaria solani* infected susceptible tomato cultivar

Results of the differentially expressed genes (DEGs) recognized through the libraries constructed from control (S) vs *Alternaria solani* (S) infected tomato leaves transcripts were presented in **Fig.2**. In total, 23670 transcripts were found to be expressed, out of which 9,864 were significantly expressed. A total of 1,248 were found to be upregulated, and 1,562 were downregulated. There were 105 and 58 transcripts specific to Control (S) and tomato leaves infected with *Alternaria solani* alone in susceptible cultivars.

The differentially expressed genes that were obtained between control and *A. solani* infected susceptible tomato cultivar were analyzed, wherein the gene that exhibited increased expression level in the susceptible tomato cultivar belonging to the control group was enzyme peptidylprolyl isomerase, which is involved in protein folding and unfolding as well as in protein transport (Zgajnar *et al*., 2019). However, a few genes, including Cytochrome P450 and Cytochrome P450-dependent fatty acid hydroxylase, were found to be downregulated in susceptible control tomato cultivars. Studies have shown that in grapevine, P450 protects the plant from biotic and abiotic stresses (Minerdi *et al*., 2023), which substantiates how a higher level of P450 is found in *A. solani* infected tomato plants compared to control.

Interestingly, higher expression of gene Ethylene-responsive proteinase inhibitor 1 was observed only in *A. solani* infected leaves of susceptible cultivar of tomato, which was absent in the control. Diaz *et al*. (2002) revealed that tomato plants were shown to have higher levels of proteinase inhibitor 1 when tomato plants were infected with *Botrytis cinerea.* Chlorophyll a-b binding protein and Class II, small heat shock proteins, were upregulated in a susceptible tomato cultivar infected with A. solani.In *Triticum aestivum*, light-harvesting chlorophyll a/b binding protein, TaLhc2 induced the ramification of phytopathogenic fungus and, as a result, showed a higher level of PR protein expression (Han *et al*., 2023). Similarly, overexpression of OsHsp18.0, a class II small heat shock protein gene in a susceptible rice cultivar, significantly enhanced its resistance to *Xanthomonas oryzae* pv. *oryzae* (Kuang *et al*., 2017). One of the significantly downregulated genes was the protein in the BTB domain. Furthermore, CaBPM4, a BTB domain-containing gene, was downregulated in *Capsicum annuum* L. upon infection with *Phytophthora capsici* (He *et al*., 2019).

**Fig. 2.**
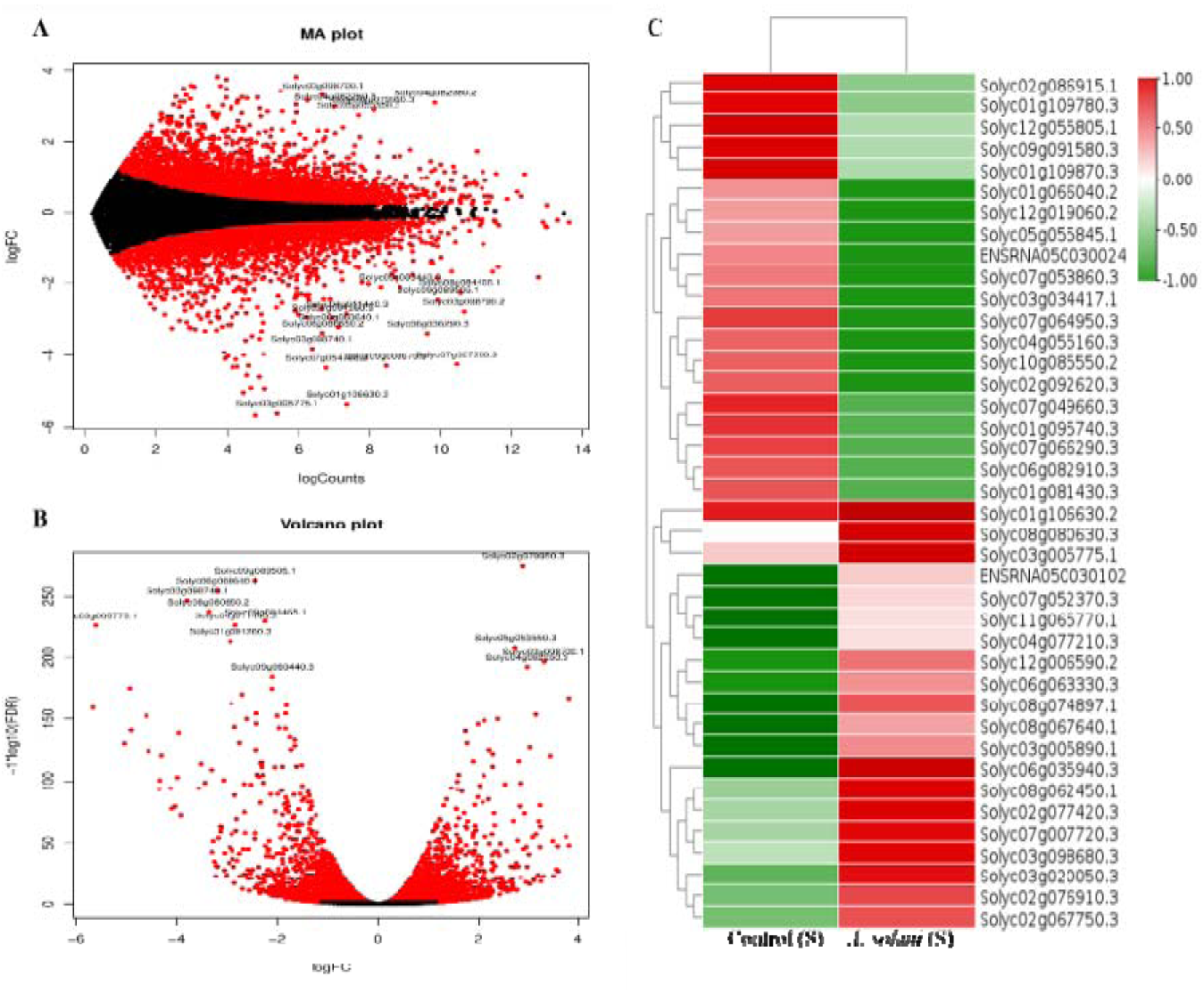
Transcriptome analysis of differentially expressed genes (DEGs) of control (S) *vs A. solani* (S) infected *S. lycopersicum.* **A.** MA plot and **B.** Volcano plot shows red and black colour dots showed the significantly expressed upregulated and downregulated genes respectively. **C.** Heat map and hierarchical clustering represent the DEGs. Red color shows the upregulated genes, and green shows the downregulated genes (S - Susceptible cultivar)

#### Comparison of differentially expressed genes between control and *A. solani* infected tolerant tomato cultivar

The analysis of the DEGs in the uninfected tolerant tomato cultivar and tolerant tomato cultivar infected with *A. solani* provided 23259 transcripts, out of which 14,691 were significantly expressed. From the significantly expressed genes, 4,030 transcripts were upregulated, and 4,544 transcripts were downregulated. In **Fig.3**, one of the upregulated genes in tolerant control tomato cultivar was Glycosyltransferase, wherein the overexpression of *the WssgtL3.1* gene revealed less infestation of *Pseudomonas syringae* in *Arabidopsis thaliana* due to the accumulation of Salicylic acid (Mishra *et al*., 2022). Dehydrins and Protein LE25, II late embryogenesis abundant (LEA) proteins, were downregulated in tolerant control tomato cultivar. The gene expression of dehydrin was comprehensively studied through the regulatory effects of ABA, MAPK, and second messenger calcium by Sun *et al*., 2021.

Further, the dataset also showed the downregulation of AAA+ ATPase domain-containing protein in tomato plants (Tolerant cultivar) infected with *A. solani*. *LMR* encodes an AAA-type ATPase that is localized to the chloroplast. The LMR gene has been reported to provide information about elucidating molecular mechanisms underlying PCD in plants. However, the defense response involving PCD is less understood in monocots compared with the mechanisms in dicots (Matin *et al*., 2010). In *Nicotiana benthamiana*, the yeast two-hybrid system-based studies indicated that these proteins interact with the helicases of the TMV replicase enzyme, which contains a conserved AAA family motif (Abbink *et al*., 2002). Interestingly, a tolerant cultivar of tomato plants protects itself from *A. solani* infection by downregulating the AAA+ ATPase domain-containing protein.

Heat map and hierarchical clustering indicate that the non-specific lipid transfer proteins (LTP) were upregulated in uninfected and in *A. solani* pathogen alone infected tolerant tomato cultivar. Several overexpression studies have reported that nsLTPs could resist diseases caused by plant pathogens. For example, Wang *et al*. (2021) studied the overexpression of StLTP10, a non-specific lipid transfer protein, in *Solanum tuberosum* and found that the gene showed higher resistance against *Phytophthora infestans*. Similarly, Bvindi *et al*. 2023 overexpressed a non-specific lipid transfer protein, StnsLTP1.33, in *Solanum tuberosum* and conferred enhanced susceptibility to *Alternaria solani*. In addition, suppression of CaLTPs by virus-induced gene silencing (VIGS) proved enhanced susceptibility to *Xanthomonas campestris* pv. *vescatoria* and pepper mosaic mottle virus in pepper (Sarowar *et al*., 2019). Thus, the present study suggests that the upregulated genes in uninfected and infected tolerant tomato cultivar could be essential candidate genes for susceptibility or resistance factors.

**Fig. 3.**
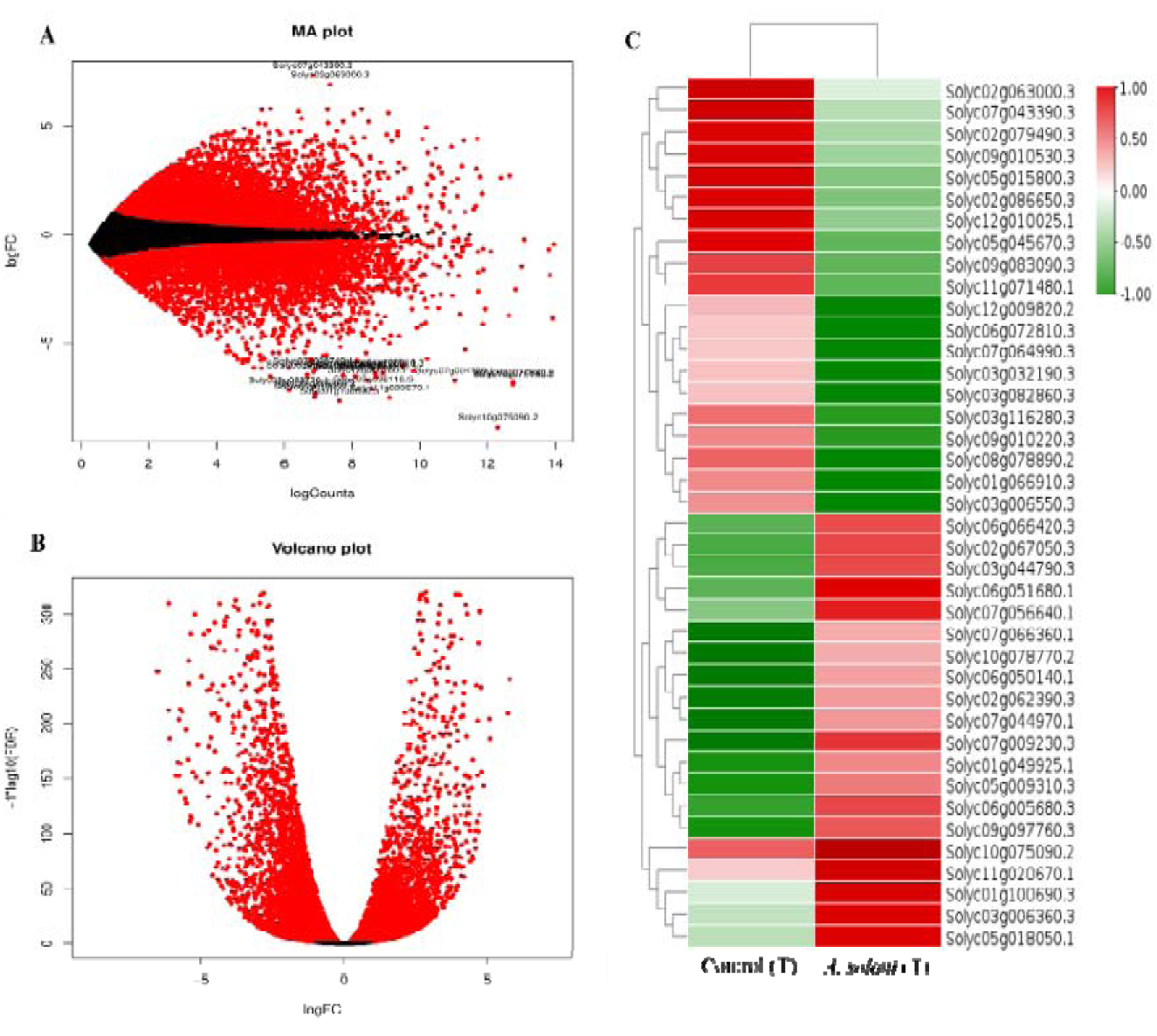
Transcriptome analysis of differentially expressed genes (DEGs) of control (T) *vs A. solani* (T) infected with *S. lycopersicum.* **A.** MA plot and **B.** Volcano plot represented in red and black color dots showed the significantly expressed upregulated and downregulated genes, respectively. **C.** Heat map and hierarchical clustering represent th DEGs. The red color showed the upregulated genes, and the green color showed th downregulated genes (T - Tolerant cultivar).

#### Comparison of differentially expressed genes between tolerant and susceptible tomato cultivar upon *A. solani* infection

While comparing the DEGs between tolerant tomato cultivars as well as susceptible tomato cultivars infected with *the A. solani* pathogen, 23108 transcripts were found to be expressed, out of which 13848 were significantly expressed. A total of 3796 and 3919 were found to be upregulated and downregulated, respectively, from which 323 and 181 transcripts were specific to *Solanum lycopersicum* leaves infested with *A. solani* in susceptible and tolerant cultivars. Interestingly, Heat shock protein 70, Heat shock factor (HSF) type DNA binding domain-containing protein, and Co-chaperone protein p23 were upregulated in susceptible tomato cultivars infected with *Alternaria solani* pathogen only (**Fig.4**.). Studies have shown that Heat shock proteins play an essential role in defense response against biotic stresses. Kim & Hwang (2015) studied the silencing of the HSP70a gene in *Capsicum annum*, which promoted the manifestation of *Xanthomonas campestris* pv *vesicatoria* and reduced the defense marker gene expression. Studies revealed that p23 is one of the essential molecular co-chaperones of Hsp90 (Zhang *et al*., 2010)

Inositol-3-phosphate synthase was downregulated in susceptible tomato cultivars infected with *the Alternaria solani* pathogen. However, a slight upregulation was observed in tolerant tomato cultivars infected with the *Alternaria solani* pathogen. In Arabidopsis thaliana, the plants mutated with the Inositol-3-phosphate synthase gene were hypersusceptible to tobacco mosaic virus, turnip mosaic virus, cucumber mosaic virus, and cauliflower mosaic virus as well as to the fungus *Botrytis cinerea* and *Pseudomonas syringae* (Murphy *et al*., 2018). Non-specific lipid-transfer protein was upregulated in susceptible and tolerant tomato cultivar infected with *Alternaria solani.* DUF4005 domain-containing protein, RING-type domain-containing protein, and Protein early flowering 4 domain-containing protein were a few genes that were upregulated in tolerant tomato cultivars infected with *Alternaria solani*. B box-type domain-containing protein is a downregulated gene in tolerant tomato cultivars infected with *Alternaria solani*. Alternatively, the B box-type domain-containing gene, *IbBBX24* was found to be involved in the regulation of the jasmonic acid pathway in *Ipomoea batatas*, and overexpression of *IbBBX24* gene induced the disease resistance against *Fusarium oxysporum* f. sp *batatas (*Zhang *et al*., 2020).

**Fig. 4.**
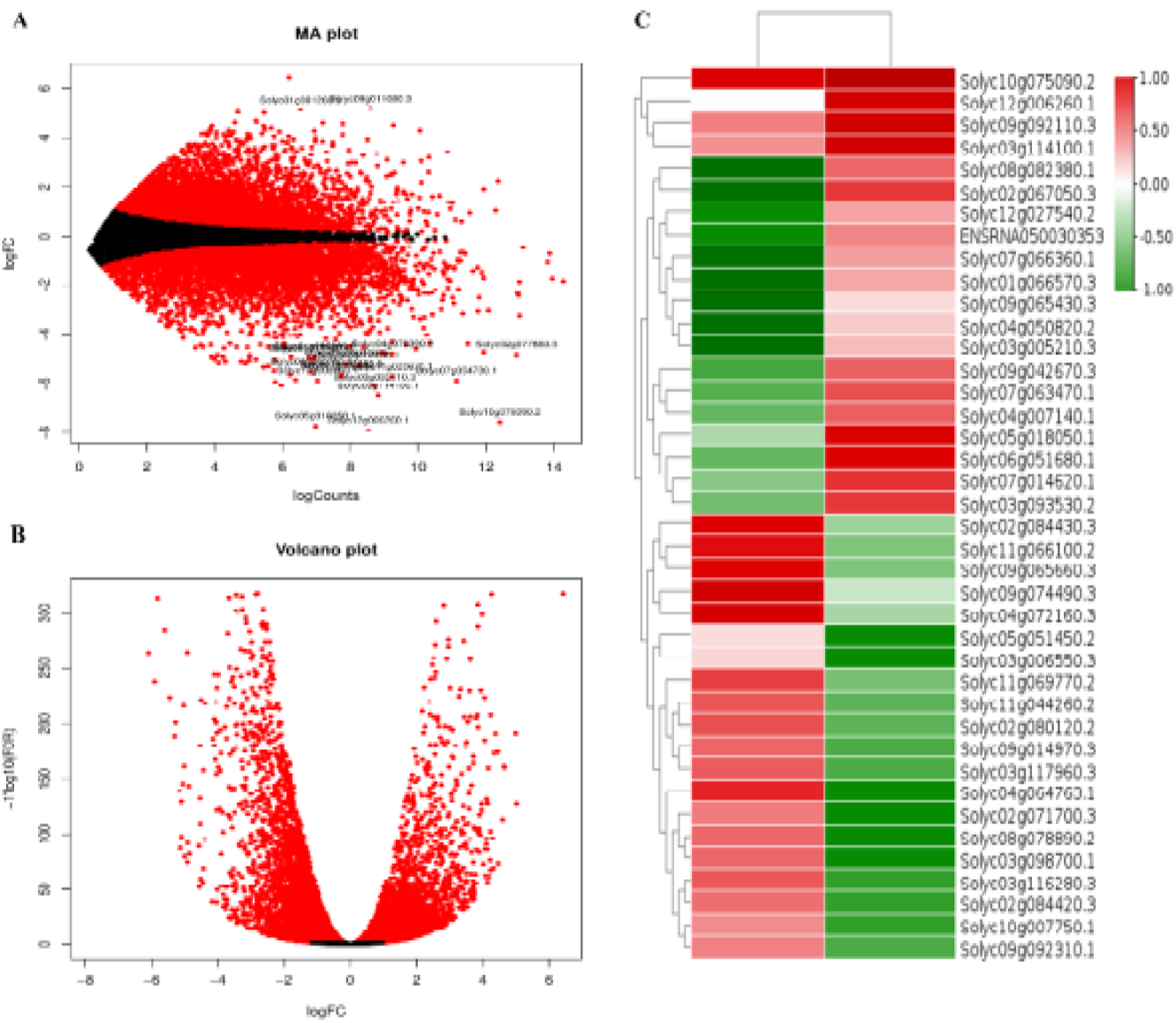
Transcriptome analysis of differentially expressed genes (DEGs) of *A. solani* (S) *vs A. solani* (T) infected with *S. lycopersicum.* **A.** MA plot and **B.** Volcano plot indicated in red and black color dots showed the significantly expressed upregulated and downregulated genes, respectively. **C.** Heat map and hierarchical clustering represent the DEGs. Red color showed the upregulated genes, and green colors showed the downregulated gene (S - Susceptible and T - Tolerant cultivars).

### Transcriptional changes in tolerant and susceptible tomato cultivar upon Laminarin pretreatment

#### Comparison of differentially expressed genes between control and LaM pretreated susceptible tomato cultivar

A significant upregulation of genes was observed in the elicitor-treated samples compared to the control of susceptible tomato cultivars. A total of 23537 transcripts were found to be expressed, out of which 9996 were significantly expressed. About 1146 and 1279 gene were found to be upregulated and downregulated, respectively. In susceptible cultivar, 85 and 31 transcripts were specific control (S) and Laminarin (S). In the untreated control plants, as well as in the susceptible tomato cultivar pretreated with the elicitor laminarin, the tetratricopeptide repeat protein showed a significant increase in both plants (**Fig.5**.). According to Rosado et al. (2006), tetratricopeptide repeat proteins regulated several dehydration-responsive proteins such as *ERD1*, *ERD3*, *and COR15a*. Zhang et al. (2022) discovered that *BnaA02.YTG1* encodes a chloroplast-localized tetratricopeptide repeat protein expressed widely in *Brassica napus* and is essential for early chloroplast biogenesis.

In control tomato plants belonging to susceptible cultivars, EF-hand domain-containing protein, which is a calcium ion binding protein, as well as Reverse transcriptase/retrotransposon-derived protein RNase H-like domain-containing protein, a reverse transcriptase enzyme, were found to be upregulated. Few genes, such as LRAT domain-containing protein, Premnaspirodiene oxygenase-like, RRM domain-containing protein, and Fe2OG dioxygenase domain-containing protein, were found to be downregulated in control tomato plants belonging to susceptible cultivars when compared to Laminarin pretreated susceptible tomato cultivar. For instance, Premnaspirodiene oxygenase-like protein is a cytochrome P450 enzyme that catalyzes the hydroxylation reactions of different sesquiterpene substrates (Takahashi *et al*., 2007). Similarly, Zabel *et al*. (2021) also revealed that zingiberene oxidase, produced from *Solanum habrochaites,* shown to undergo sequential oxidations like Premnaspirodiene oxygenase-like protein, which is a sesquiterpene oxidase. In addition, Yang *et al*. (2020) suggested that SlORRM4, an organelle RNA recognition motif (RRM) of *Solanum lycopersicum*, is required for mitochondrial function and fruit ripening. Wei *et al*. (2021) characterized several Fe2OG dioxygenases belonging to *Solanum lycopersicum* and showed that they were involved in metabolic pathways like gibberellin biosynthesis, ethylene biosynthesis, steroidal glycoalkaloid biosynthesis, and flavonoid metabolism.

**Fig. 5.**
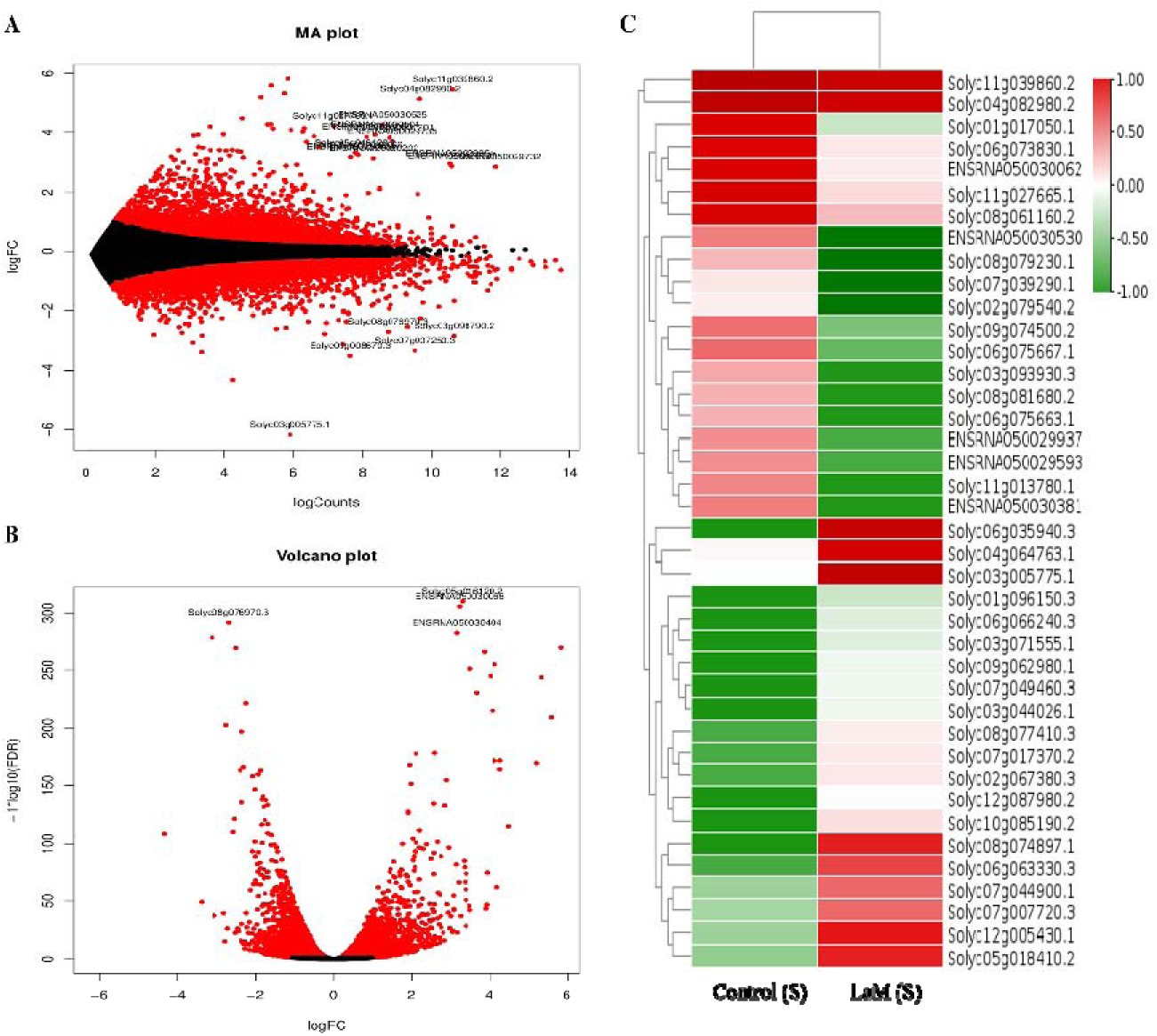
Transcriptome analysis of differentially expressed genes (DEGs) of control (S) *vs* Laminarin (S) pretreated with *S. lycopersicum.* **A.** MA plot and **B.** Volcano plot indicated in red and black color dots showed the significantly expressed upregulated and downregulated genes respectively. **C.** Heat map and hierarchical clustering represent the DEGs. Red color showed the upregulated genes, and green colors showed the downregulated genes (S - Susceptible cultivar).

#### Comparison of differentially expressed genes between control and LaM pretreated tolerant tomato cultivar

In total, 22774 transcripts were found to be expressed, out of which 9437 were significantly expressed. A total of 1004 and 1633 were found to be upregulated and downregulated, respectively, in **Fig. 6**. There were 101 and 18 transcripts specific control and tomato leaves pretreated with Laminarin in the tolerant cultivar. In tolerant cultivars, Cytosol aminopeptidase domain-containing protein, Enoyl reductase (ER) domain-containing protein, and Peptidase M20 dimerization domain-containing protein were upregulated in control tomato plants. Interestingly, Cytosol aminopeptidase domain-containing protein was upregulated in tolerant cultivars of both control and laminarin-pretreated tomato plants. Studies have shown that tomato leucine aminopeptidases (LAPs) have dual locations in the Cytosol as well as in plastids, in which LAP-A proteins are primarily detected within chloroplast (Narváez-Vásquez *et al*., 2008).

The genes that are upregulated in control (T) were Cytosol aminopeptidase domain-containing protein, TPSI1 protein, Enoyl reductase (ER) domain-containing protein, Peptidase M20 dimerization domain-containing protein and K-box domain-containing protein. The genes which are downregulated were Na+ transporter, Chalcone synthase and Nucleotide-diphosphate-sugar transferase domain-containing protein, VQ domain-containing protein, and Bifunctional inhibitor/plant lipid transfer protein/seed storage helical domain-containing protein. Genes explicitly upregulated in Laminarin (T) pretreated with tomato plants included TPSI1 protein, AP2/ERF domain-containing protein, EF-hand domain-containing protein, NADP-dependent oxidoreductase domain-containing protein, and Methylesterase. Genes downregulated in Laminarin (T) pretreated with tomato plants were K-box domain-containing protein, RNase III domain-containing protein, Fe2OG dioxygenase domain-containing protein, and RRM domain-containing protein. Interestingly, upregulation of TPSI1 protein was observed in both control, and Laminarin pretreated tolerant tomato cultivars. In *Arabidopsis thaliana*, *AtIPS1* belongs to the *TPSI1/Mt4* family, the members of which are specifically induced by Pi starvation and, as a result, Pi deprivation-induced *AtIPS1* expression in all cells of wild-type plants (Martin *et al*., 2000).

**Fig. 6.**
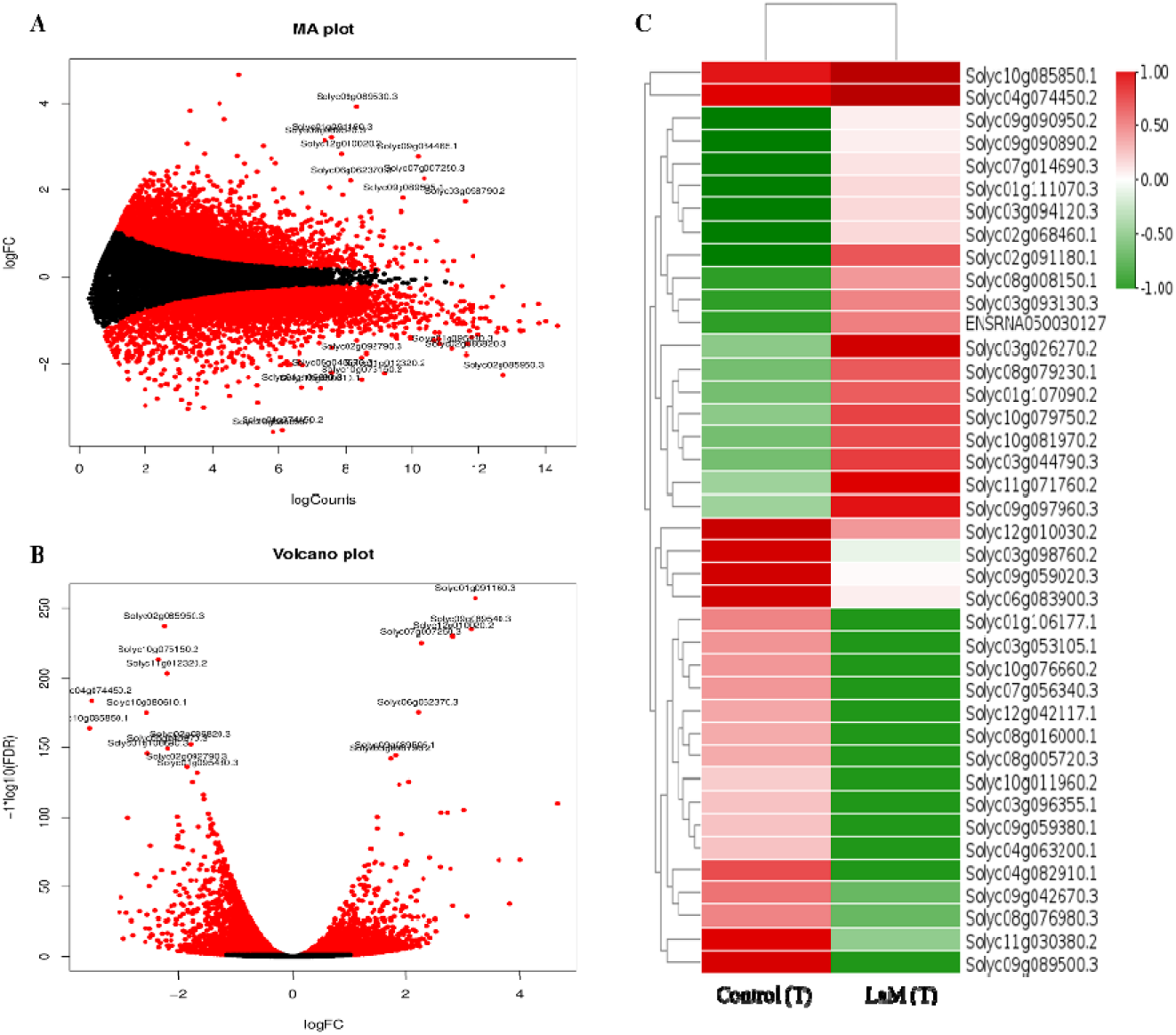
Transcriptome analysis of differentially expressed genes (DEGs) of control (T) *vs* Laminarin (T) pretreated with *S. lycopersicum.* **A.** MA plot and **B.** Volcano plot represented in red and black color dots showed the significantly expressed upregulated and downregulated genes, respectively. **C.** Heat map and hierarchical clustering represent the DEGs. Red color showed the upregulated genes, and green colors showed the downregulated genes (Tolerant Cultivar)

#### Comparison of differentially expressed genes between LaM pretreated tolerant and susceptible tomato cultivar

Results of the differentially expressed genes (DEGs) recognized through the libraries constructed from Laminarin (S) *vs* laminarin (T) laminarin pretreated with tomato resulted in a total of 10251 transcripts. In total, 22713 transcripts were found to be expressed, out of which 10251 were significantly expressed. 1156 and 1837 were found to be upregulated and downregulated, respectively. There were 209 and 50 transcripts specific to tomato leaves pretreated with Laminarin in susceptible and tolerant cultivars (**Fig.7**.).

The genes upregulated in Laminarin (S) were: C3H1-type domain-containing protein, RNase III domain-containing protein, Peptidase M20 dimerization domain-containing protein, DNA-directed RNA polymerase, and ADP-ribosylation factor. The genes which are downregulated were OTU domain-containing proteins), Transmembrane protein, Chalcone synthase, Protein FAR1-Related sequence, and F-box domain-containing protein. Genes that are explicitly upregulated in Laminarin (T) alone pretreated with tomato plants were Sugar phosphate transporter domain-containing protein, AP2/ERF domain-containing protein, Aquaporin (Plasma membrane intrinsic protein 24), nicotianamine synthase and Nuclear pore complex protein NUP88. Genes that were downregulated in Laminarin (T) pretreated with tomato plants were Pentatricopeptide repeat-containing protein, Glycosyltransferase, Bet v I/Major latex protein domain-containing protein, and Mediator complex subunit 15 KIX domain-containing protein.

**Fig. 7.**
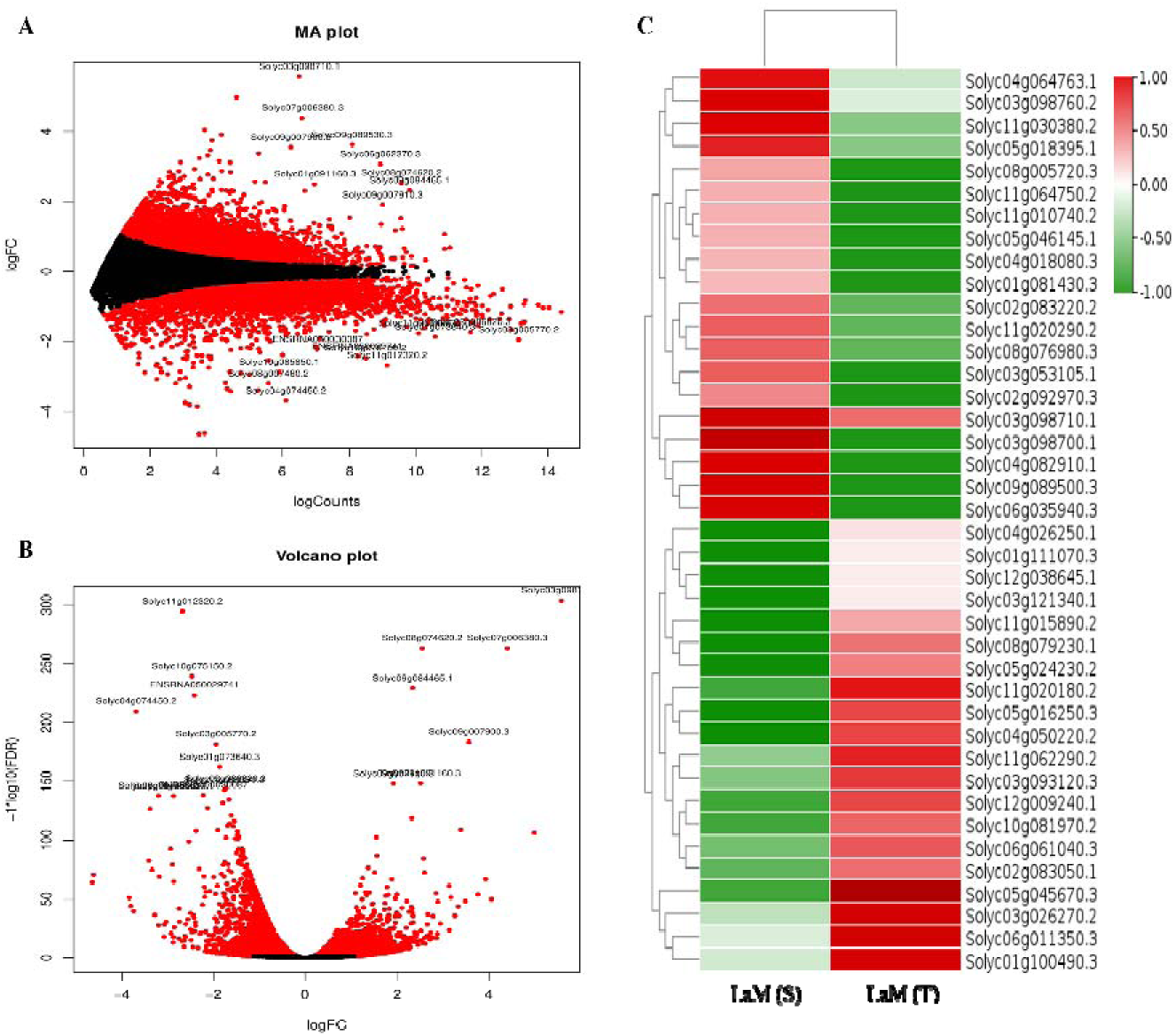
Transcriptome analysis of differentially expressed genes (DEGs) of Laminarin (S) *vs* Laminarin (T) pretreated with *S. lycopersicum.* **A.** MA plot and **B.** Volcano plot, show the significantly expressed upregulated and downregulated genes, respectively. **C.** Heat map and hierarchical clustering represent the DEGs. Red color showed the upregulated genes, and green colors showed the downregulated genes (S - Susceptible and T - Tolerant cultivars).

### Transcriptional changes in tolerant and susceptible tomato cultivar upon Laminarin pretreatment followed by *Alternaria solani* infection

#### Comparison of differentially expressed genes between LaM pretreated and LaM pretreated susceptible tomato cultivar followed by *Alternaria solani* infection

In susceptible tomato cultivars pretreated with Laminarin and Laminarin followed by *A. solani* infection, 22971 transcripts were found to be expressed, out of which 12031 were significantly expressed. A total of 2108 and 2500 were found to be upregulated and downregulated, respectively. There were 94 and 46 transcripts specific laminarin (S) and Laminarin *+ A. solani (S)* tomato leaves pretreated with Laminarin and/or infected with *A. solani* in susceptible cultivars (**Fig. 8**.).

The susceptible tomato cultivar pretreated with Laminarin showed significant upregulation of genes such as Bifunctional inhibitor/plant lipid transfer protein/seed storage helical domain-containing protein, Fructose-bisphosphatase, Protein kinase domain-containing protein, DUF4005 domain-containing protein and Zinc finger CCCH domain-containing protein 18. Similarly, Aquaporin Plasma membrane intrinsic protein 24 (PIP) was significantly upregulated in susceptible cultivars of tomato plants pretreated with Laminarin and infected by *A. solani*. The oxidative burst is a vital defense response of plants to pathogen infection resulting in the generation of ROS around the site of infection (Pitzschke *et al*., 2009; Wang *et al*., 2019). The rapid and transient generation of ROS H_2_O_2_ indicates explicitly successful recognition of pathogen invasion. This initiates defense responses, such as PTI and SAR, to regulate plant disease resistance (Torres, 2010). It has been reported that AtPIP1;4, a member of the PIP family in *Arabidopsis*, was documented in the transport of H_2_O_2_ through the PM (Liu & He, 2016). It activates MAPK signaling by expressing *MPK3*, producing Callose, and regulating SAR’s *NPR1* and *PR* genes (Liu & He, 2016). *AtPIP1;4* knockout mutants study also revealed that H_2_O_2_ was prevented from being transported into the cells, and high concentrations of H_2_O_2_ accumulated within apoplastic regions, leading to hypersensitivity to the pathogen. These findings indicated the role of Aquaporin Plasma membrane intrinsic protein in translocating apoplastic H_2_O_2_, leading to the activation of the PTI and SAR pathways in response to PAMPs.

Genes that were downregulated in Laminarin + *A. solani* (S) pretreated with tomato plants were Non-specific serine/threonine protein kinase, Terpene synthase N-terminal domain-containing protein, Pentatricopeptide repeat-containing protein, Pentacotripeptide-repeat region of PRORP domain-containing protein. Interestingly, higher expression of Bifunctional inhibitor/plant lipid transfer protein/seed storage helical domain-containing protein was observed in Laminarin alone pretreated susceptible tomato cultivar as well as in laminarin pretreatment followed by *A. solani* (S) infected susceptible tomato cultivar (**Fig. 7**).

**Fig. 8.**
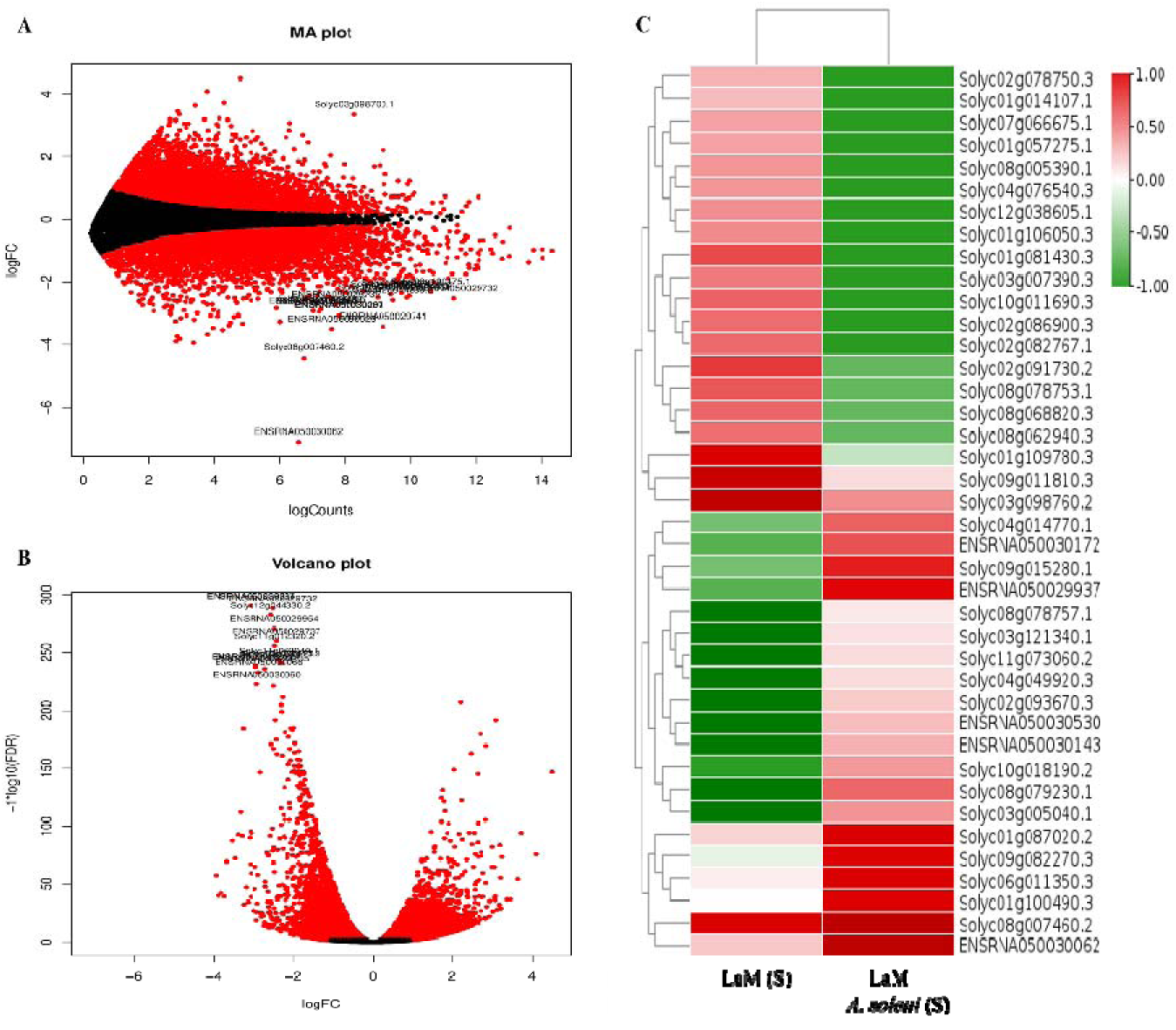
Transcriptome analysis of differentially expressed genes (DEGs) of Laminarin (S) *vs*. laminarin + *A. solani* (S) pretreated with *S. lycopersicum.* **A.** MA plot and **B.** Volcano plot shows the significantly expressed upregulated and downregulated gene respectively. **C.** Heat map and hierarchical clustering represent the DEGs. Red color showed the upregulated genes, and green colors showed the downregulated genes (S - Susceptible cultivar).

#### Comparison of differentially expressed genes between LaM pretreated and LaM pretreated tolerant tomato cultivar followed by *Alternaria solani* infection

Out of 22711 transcripts, 10909 genes were differentially expressed in libraries constructed from tolerant tomato cultivar pretreated with Laminarin vs. tolerant tomato cultivar pretreated with Laminarin followed by *A. solani* infection. A total of 2421 and 1581 were found to be upregulated and downregulated, respectively; 15 genes were specifically expressed in a tolerant cultivar of tomato pretreated with Laminarin compared to 201 genes that were significantly expressed during pretreatment of Laminarin followed by *A. solani* infection in a tolerant cultivar of tomato.

Flower-specific gamma-thionin-like protein/acidic protein, Reverse transcriptase domain-containing protein, AP2/ERF domain-containing protein and DUF4228 domain-containing protein were some of the genes that were upregulated in tolerant tomato cultivar pretreated with Laminarin alone, while Pectate lyase, Peroxidase, HMA domain-containing protein, RRM domain-containing protein and Kinesin motor domain-containing protein were few downregulated genes.

There was a significant upregulation of the Sugar phosphate transporter domain-containing protein in tomato plants pretreated with Laminarin, followed by *A. solani* infection in tolerant cultivars of tomato. Interestingly, in Arabidopsis leaves challenged with *Botrytis cinerea*, increased expression of STP13 and stp13 mutant plants exhibit enhanced susceptibility and reduced glucose uptake rates. In contrast, STP13 overexpressing plants show a resistant phenotype and higher glucose transport capacity involved in the active resorption of hexoses from the apoplast, depriving the pathogen of its sugar source, thereby reducing the pathogen infection (Lemonnier *et al*., 2014; Breia *et al*., 2021).

Specific genes that were downregulated in tolerant tomato cultivar pretreated with Laminarin followed by *A. solani* infection were Reverse transcriptase domain-containing protein, Protein kinase domain-containing protein, VQ domain-containing protein, Protein transport protein SEC23, and Serine-threonine/tyrosine-protein kinase catalytic domain-containing protein. Interestingly, higher expression of an uncharacterized protein (Solyc09g059020.3) was observed only in tolerant tomato cultivar pretreated with Laminarin followed by *A. solani* infection compared with Laminarin alone pretreated tolerant tomato cultivar (**Fig. 9**.).

**Fig. 9.**
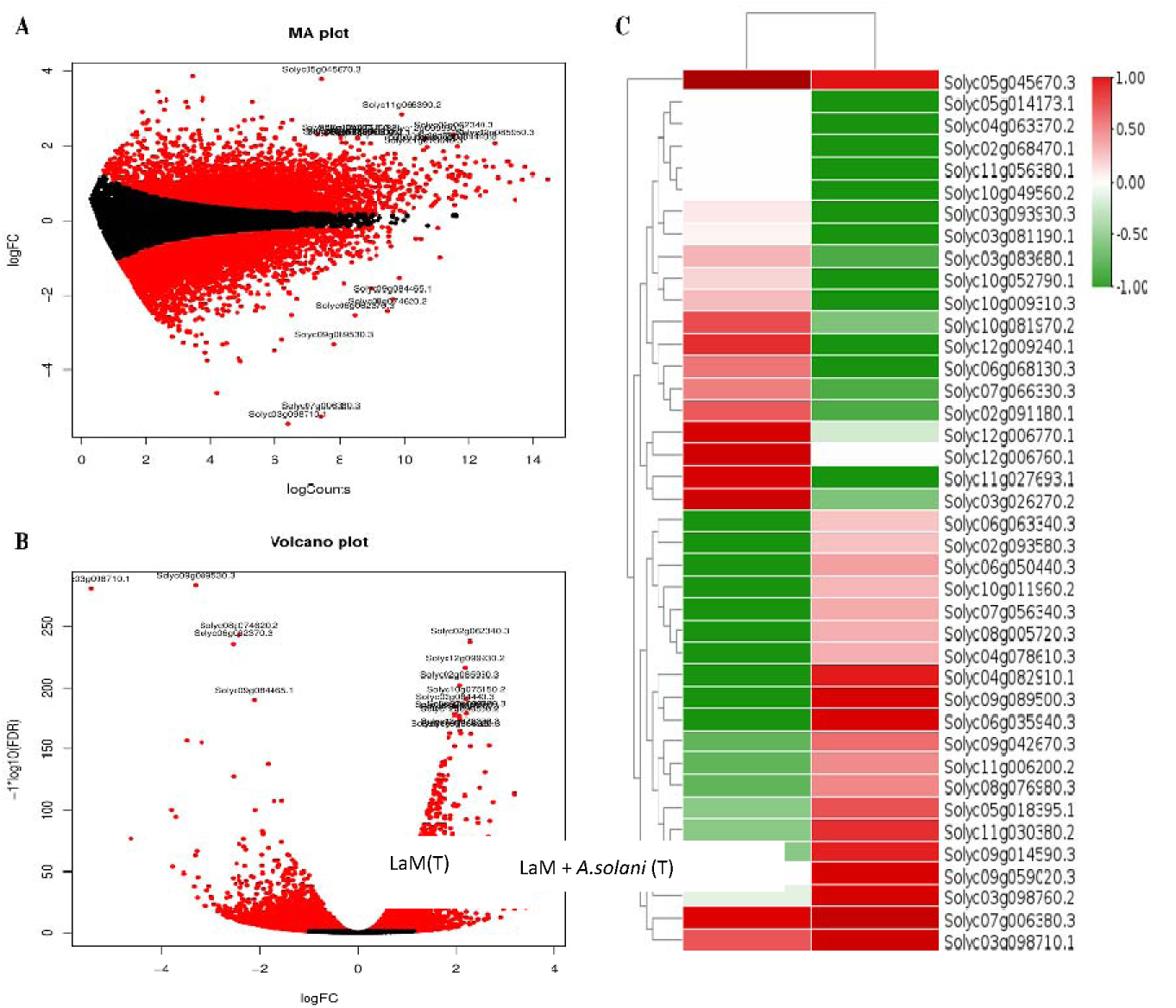
Transcriptome analysis of differentially expressed genes (DEGs) of Laminarin (T) *vs* laminarin + *A. solani* (T) pretreated with *S. lycopersicum.* **A.** MA plot and **B.** Volcano plot shows red and black color dots showed the significantly expressed upregulated and downregulated genes respectively. **C.** Heat map and hierarchical clustering represent the DEGs. Red color showed the upregulated genes, and green colors showed the downregulated genes (T - Tolerant cultivar).

#### Comparison of differentially expressed genes between LaM pretreated susceptible tomato cultivar followed by *Alternaria solani* infection, and LaM pretreated tolerant tomato cultivar followed by *Alternaria solani* infection

Experimental tomato leaves of susceptible and resistant cultivars were treated with elicitor LaM followed by *A. solani* pathogen inoculation. RNA seq analysis showed 23097 DEGs, including 12709 significant transcripts, with 2845 upregulated genes. According to th heat map and hierarchical clustering, the DEGs that showed higher expression were Elongation factor 1 alpha, Peptidase 554 rhomboid domain-containing protein, Major facilitator superfamily profile domain-containing protein, Mitochondrial pyruvate carrier, Reverse transcriptase domain-containing protein (**Fig. 10**.). A total of 2297 genes were downregulated, including Timeless N-terminal domain-containing protein, Protein kinase domain-containing protein, Kinesin motor domain-containing protein, Pentatricopeptide-repeat region of PROPO domain-containing protein, and AAA+ ATPase domain-containing protein.

DUF 4005 domain-containing protein, sugar phosphate transporter domain-containing protein, meiotic recombination protein DMC1 homolog, Leucine-rich repeat-containing N-terminal plant type domain-containing protein, Subtilisin like protease were explicitly upregulated in tomato plants pretreated with LaM followed by *A. solani* (T). In plants, leucine-rich repeat (LRR) regions are responsible for the induction of signal transduction pathways upon pathogen infection, ultimately activating defense-related genes. Though LRRs have no direct inhibitory effect against pathogens, these repeats in resistance genes enable them to recognize avirulence gene products (avr) of the pathogen and to induce corresponding responses in host plants. Extracytoplasmic LRR motifs also play a significant role for specific defense proteins, such as polygalacturonase inhibiting proteins (PGIPs) and extensions (Jones & Jones, 1997). However, they are not directly involved in pathogen recognition and activation of defense genes (Stotz *et al*., 2000). The limitation of hyphal intrusion by PGIPs also activates the expression of defense components in plants. However, without specific recognition at the PG-PGIP level, the pathogen evades plant defense. (Sharrock and Lubavitch, 1994).

Genes that were downregulated explicitly in tomato plants pretreated with LaM followed by *A. solani* (T) were Reverse transcriptase domain-containing protein, NADH: quinone oxidoreductase/Mrp antiporter membrane subunit domain-containing protein, SWIM-type domain-containing protein, Peroxisomal membrane protein PMP22, and AAA+ ATPase domain-containing protein. However, higher expression of Peptidase 554 rhomboid domain-containing protein was observed only in plants treated with Laminarin followed by *A. solani* susceptible cultivar, while tolerant tomato cultivar treated with Laminarin followed by *A. solani* infection did not show significant gene expression.

**Fig. 10.**
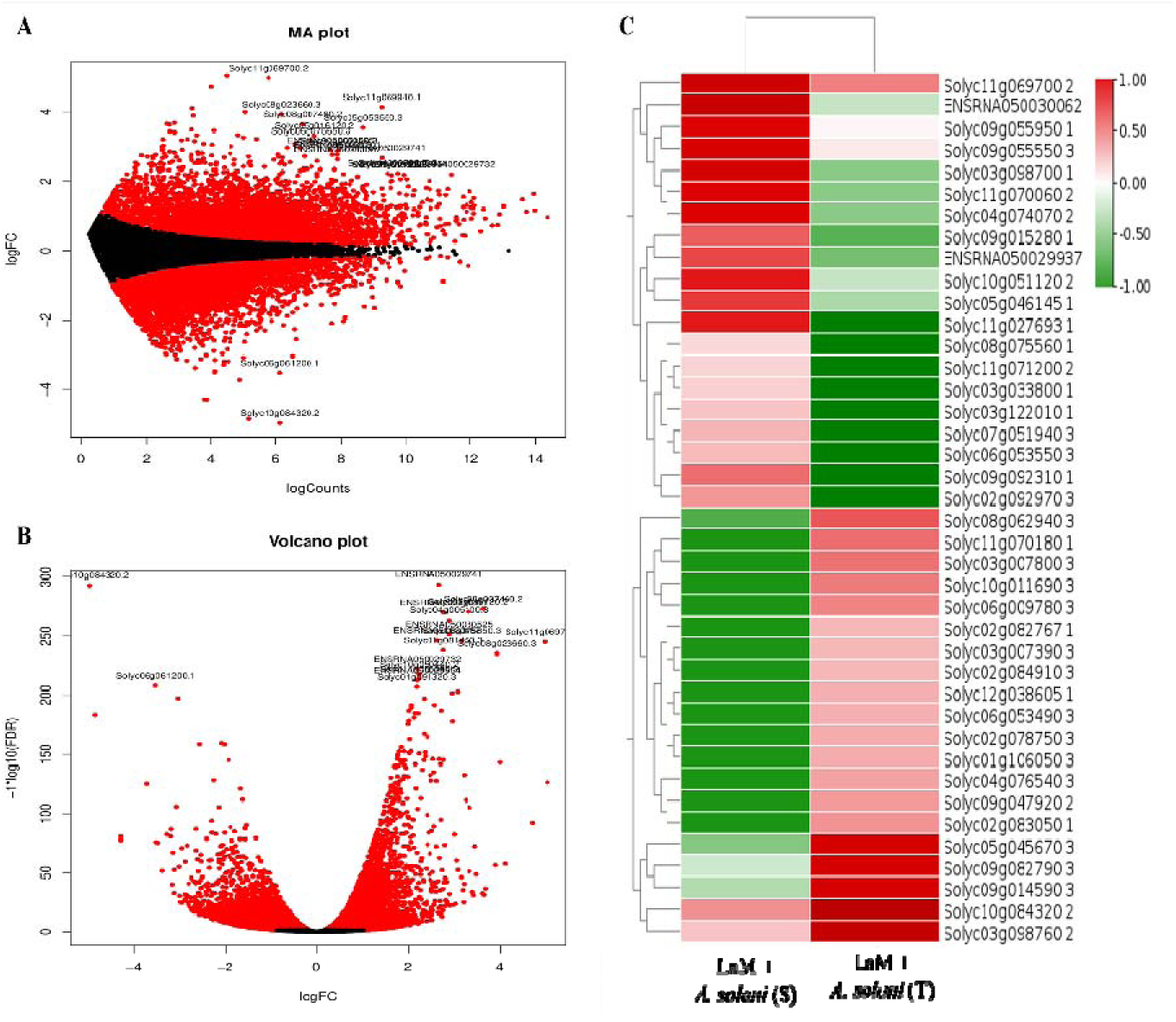
Transcriptome analysis of differentially expressed genes (DEGs) of laminarin + *A. solani* (S) *vs* Laminarin + *A. solani* (T) pretreated with *S. lycopersicum.* **A.** MA plot and **B.** Volcano plot shows red and black color dots showed the significantly expressed upregulated and downregulated genes respectively. **C.** Heat map and hierarchical clustering represent the DEGs. The red color showed the upregulated genes, and th green showed the downregulated genes (S & T - Susceptible & Tolerant cultivars).

#### Gene Ontology Enrichment Analysis & Pathway Enrichment of the differentially expressed genes in control and *A. solani* infected susceptible tomato cultivar

Results of gene ontology (GO) enrichment analysis show enrichment of differentially expressed genes in biological processes, cellular components, and molecular functions (**S. Fig. 1. A**). In the differentially expressed genes obtained between control and *A. solani* infected susceptible tomato cultivar, maximum gene expression in biological processes belongs to protein phosphorylation (378 genes) and regulation of DNA-template transcription (248 genes). A minimum gene expression belongs to phosphate ion transport (98). In the case of the cellular process, genes confined to the membrane (1685) showed higher expression compared to another category of genes expressed. In the case of molecular functions, genes corresponding to ATP binding (903) showed maximum expression compared to genes involved in mRNA binding (146). In Pathway Enrichment analysis, maximum genes were involved in the protein modification pathway, whereas fewer genes were involved in porphyrin-containing compound metabolism (**S. Fig. 1. B**).

#### Gene Ontology Enrichment Analysis & Pathway Enrichment of the differentially expressed genes in control and *A. solani* infected tolerant tomato cultivar

The differentially expressed genes between control tolerant tomato cultivar and tolerant tomato cultivar infected with *A. solani* showed maximum gene expression in biological processes belonging to protein phosphorylation (561 genes) and regulation of DNA template transcription (388 genes) (**S. Fig. 2. A**). A minimum gene expression belongs to the phosphate ion transport category (142). In the case of the cellular process, genes confined to the membrane (2543) showed higher expression than the extracellular region (158), which showed minimum expression. In the case of molecular functions, genes corresponding to ATP binding (1327) showed maximum expression compared to genes involved in mRNA binding (201). In the Pathway Enrichment analysis, the maximum number of genes were involved in the protein modification pathway, whereas several genes were involved in protein glycosylation. (**S. Fig. 2. B**).

#### Gene Ontology Enrichment Analysis & Pathway Enrichment of the differentially expressed genes between *A. solani* infected susceptible and tolerant tomato cultivars

The differentially expressed genes were enriched in biological processes, cellular components, and molecular functions **(S. Fig. 3. A)**. Maximum gene expression in biological processes belongs to protein phosphorylation (517 genes) and regulation of DNA-template transcription (359 genes). A minimum gene expression is a response to a stimulus (134). In the case of the cellular process, genes confined to the membrane (2380) showed higher expression than the extracellular region (151), which showed minimum expression. In the case of molecular functions, genes corresponding to ATP binding (1238) showed maximum expression compared to genes involved in Heme binding (196). In Pathway Enrichment analysis, maximum genes were involved in the protein modification pathway, whereas fewer genes were involved in 3, 4’, and 5 – trihydroxystilbene from trans-4-coumarate (**S. Fig. 3. B**).

#### Gene Ontology Enrichment Analysis and Pathway Enrichment of the differentially expressed genes between the control and LaM-pretreated susceptible tomato cultivar

The results of gene ontology enrichment analysis reveal that expressed genes belong to biological processes, cellular components, and molecular functions (**S. Fig. 4. A)**. Maximum gene expression in biological processes belongs to protein phosphorylation (438 genes) and regulation of DNA-template transcription (231 genes). A minimum gene expression belongs to the response to the stimulus category (96). In the case of the cellular process, genes confined to the membrane (1781) showed higher expression than the extracellular region (119), which showed minimum expression. In the case of molecular functions, genes corresponding to ATP binding (971) showed maximum expression compared to genes involved in mRNA binding (149). In Pathway Enrichment analysis, the maximum number of genes were involved in the protein modification pathway, whereas several genes were involved in the 3,4’,5-trihydroxystilbene biosynthesis pathway (**S. Fig. 4. B)**.

#### Gene Ontology Enrichment Analysis and Pathway Enrichment of the differentially expressed genes between the control and the LaM-pretreated tolerant tomato cultivar

Results of gene ontology enrichment analysis reveals that genes are enriched in the category of biological processes, cellular components, and molecular functions (**S. Fig. 5. A)**. Maximum gene expression in biological processes belongs to protein phosphorylation (338 genes) and regulation of DNA-template transcription (242 genes). A minimum gene expression belongs to the defense response to the fungus category (84). In the case of the cellular process, genes confined to the membrane (1570) showed higher expression than the Golgi apparatus (107), which showed minimum expression. In the case of molecular functions, genes corresponding to ATP binding (878) showed maximum expression compared to genes involved in mRNA binding (153). In Pathway Enrichment analysis, the maximum genes were involved in the protein modification pathway, whereas several genes were involved in the Glycan metabolism pathway (**S. Fig. 5. B)**.

#### Gene Ontology Enrichment Analysis & Pathway Enrichment of the differentially expressed genes between LaM pretreated susceptible tomato cultivar and LaM pretreated tolerant tomato cultivar

The differentially expressed genes are enriched in biological processes, cellular components, and molecular functions (**S. Fig. 6. A)**. Maximum gene expression in biological processes belongs to protein phosphorylation (381 genes) and regulation of DNA-template transcription (239 genes). A minimum gene expression belongs to the defense response to fungus (94). In the case of the cellular process, genes confined to the membrane (1747) showed higher expression than the Golgi apparatus (113), which showed minimum expression. In the case of molecular functions, genes corresponding to ATP binding (964) showed maximum expression compared to genes involved in mRNA binding (158). In Pathway Enrichment analysis, the maximum genes were involved in the protein modification pathway, whereas several genes were involved in the Glycan metabolism pathway (**S. Fig. 6. B)**.

#### Gene Ontology Enrichment Analysis & Pathway Enrichment of the differentially expressed genes between LaM pretreated and LaM pretreated followed by *A. solani* infected susceptible tomato cultivar

The differentially expressed genes are enriched in biological processes, cellular components, and molecular functions (**S. Fig. 7. A)**. Maximum gene expression in biological processes belongs to protein phosphorylation (449 genes) and regulation of DNA-template transcription (290 genes). A minimum gene expression belongs to the response to the stimulus category (111). In the cellular process, genes confined to the membrane (2097) showed higher expression than the Golgi apparatus (134), which showed minimum expression. In the case of molecular functions, genes corresponding to ATP binding (1084) showed maximum expression compared to genes involved in mRNA binding (187). In Pathway Enrichment analysis, maximum genes were involved in the protein modification pathway, whereas less number of genes were involved in the Glycolysis pathway (**S. Fig. 7. B)**.

#### Gene Ontology Enrichment Analysis & Pathway Enrichment of the differentially expressed genes between the LaM-pretreated and the LaM-pretreated followed by *A. solani*-infected tolerant tomato cultivar

The differentially expressed genes are enriched in biological processes, cellular components, and molecular functions (**S. Fig. 8. A)**. Maximum gene expression in biological processes belongs to protein phosphorylation (415 genes) and regulation of DNA-template transcription (263 genes). A minimum gene expression belongs to phosphate iron transport (104). Genes confined to the membrane (1863) showed higher expression in the cellular process than the Golgi apparatus (127), which showed minimum expression. In the case of molecular functions, genes corresponding to ATP binding (1106) showed maximum expression compared to genes involved in ATP hydrolysis activity (192). In Pathway Enrichment analysis, a maximum number of genes were involved in the protein modification pathway, whereas several genes were involved in the Phytoalexin biosynthesis pathway (**S. Fig. 8. B)**.

#### Gene Ontology Enrichment Analysis & Pathway Enrichment of the differentially expressed genes between LaM pretreated followed by *A. solani* infected susceptible tomato cultivar and LaM pretreated followed by *A. solani* infected tolerant tomato cultivar

The differentially expressed genes were enriched in the biological process, cellular component, and molecular functions (**S. Fig. 9. A**). Maximum gene expression in biological processes belongs to protein phosphorylation (452 genes) and regulation of DNA-template transcription (301 genes). A minimum gene expression belongs to the carbohydrate metabolic process (116). Genes confined to the membrane (2185) showed higher expression than the Endoplasmic reticulum (139), which showed minimum expression in the cellular process. In the case of molecular functions, genes corresponding to ATP binding (1176) showed maximum expression compared to genes involved in m RNA binding (193). In Pathway Enrichment analysis, maximum genes were involved in the protein modification pathway, whereas a smaller number of genes were involved in 2-dehydro-3-deoxy-D-gluconate from the pectin pathway (**S. Fig. 9. B)**.

#### Validation of quantitative RT PCR Results

Four genes were selected for RT PCR analysis to authenticate the expression profile of RNA-seq data, and a comparison was made between the expression profile of each gene and that of RNA-seq data. This outcome implies the credibility of the transcriptome data put forth in the present study. The bZIP gene family, known as the basic leucine zipper, has been demonstrated to play a significant role in plant responses to biotic and abiotic stress. Studies on different plant species like wax gourd, bottle gourd, tobacco, and eggplant have identified numerous bZIP genes and their expression patterns under stress conditions. For instance, in wax gourd, BhbZIP58 exhibited a significant response to heat stress (Liu *et al.,* 2023).

Similarly, in tobacco, NtbZIP18 was highlighted as an essential regulator under abiotic stress conditions (Wang *et al.,* 2023). Furthermore, the SmbZIP genes in eggplants were found to be related to the control of abiotic stresses, indicating their potential role in biotic stress responses (Duan *et al*., 2022). These findings collectively emphasize the involvement of bZIP genes in plant defense mechanisms against biotic stress. The bZIP gene showed up-regulation during the *Alternaria solani* pathogen infection and LaM alone treated *Solanum lycopersicum* plants in both cultivars, which showed up to eleven-fold upregulation in the *Alternaria solani* pathogen treatment and ten-fold up-regulation in the LaM alone treatment, compared to the control and elicitor followed by pathogen treatments in the susceptible cultivar. Meanwhile, in the case of the tolerant cultivar, the minimum fold expression is shown compared to that of the susceptible cultivar. The elicitor followed by pathogen treatments in the tolerant cultivar showed an upregulation of two-fold expression compared to the control, *Alternaria solani* alone and LaM alone treatments in the susceptible cultivar (**Fig. 11**.).

**Fig. 11.**
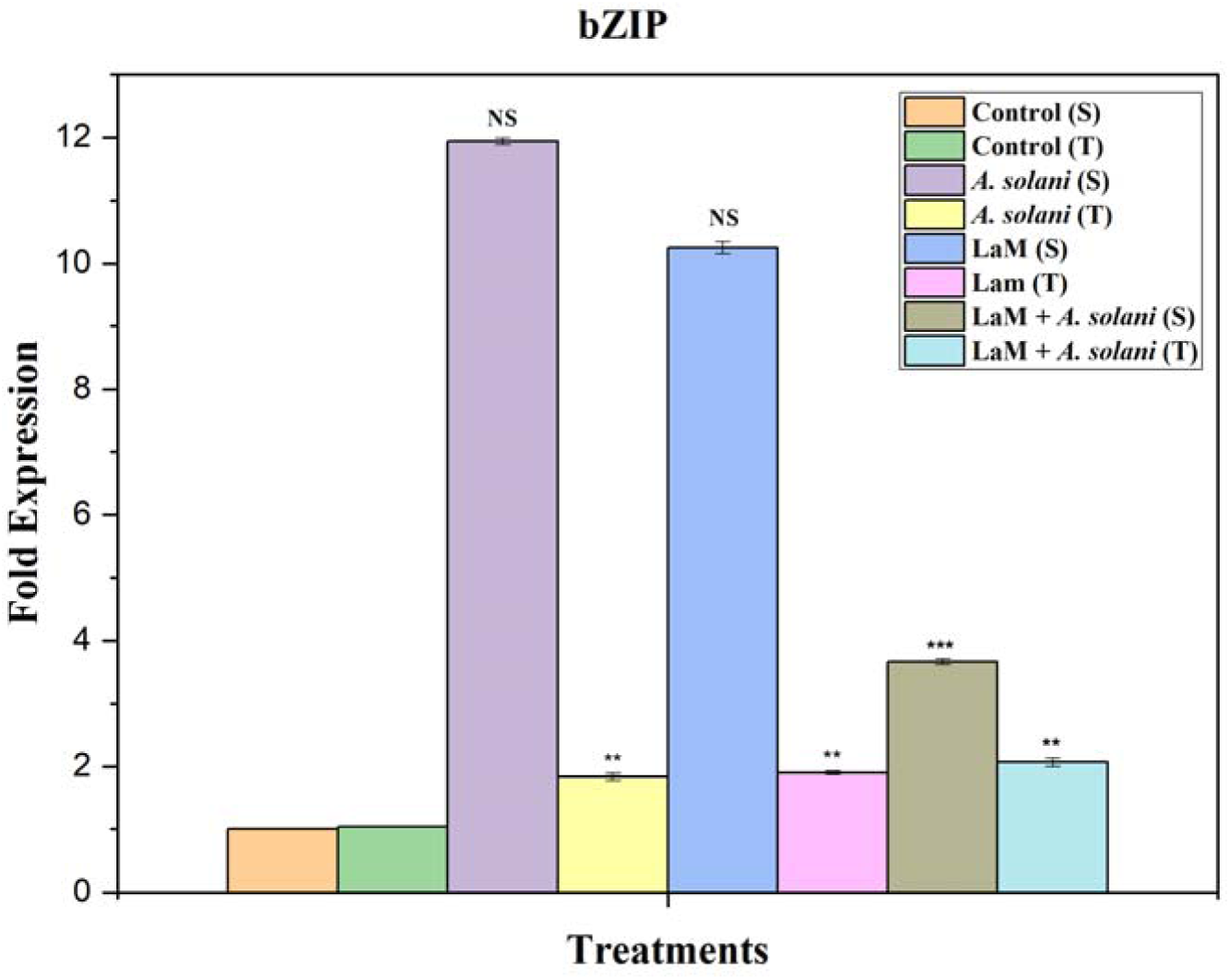
Relative expression profile of bZIP gene in Control (S), Control (T), *A. solani* (S), *A. solani* (T), LaM (S), LaM (T), LaM followed by *A. solani* (S), LaM followed by *A. solani* (T) The relative expression ratio of each gene was calculated relative to its expression in the control sample. EF1α was used as an internal control to normalize the data. Error bars representing standard deviation were calculated based on three technical replicates for three independent biological replicates. Symbols; *, **, and *** indicate significant differences between control and treated samples (i.e. p< 0.05; p< 0.01 and p< 0.001, respectively) calculated using Student’s t test in OriginPro software.

Studies have shown the involvement of Non-specific lipid transfer proteins (LTP) during biotic stress responses in *Nicotiana benthamiana*, *Cicer arietinum*, common bean, rapeseed, and Rosaceae fruit species. These genes are induced during pathogen attacks, such as tobacco mosaic virus (TMV) infection (Zhu *et al.,* 2023), *Helicoverpa armigera* infestation (Saxena *et al.,* 2023), and fungal pathogen treatments (Dong *et al.,* 2022). Expression analysis revealed differential regulation of nsLTP genes under biotic stress conditions, indicating their significance in plant defense mechanisms. The upregulation of nsLTP genes under stress conditions suggests their role in enhancing plant immunity and combating biotic challenges, emphasizing the importance of nsLTPs in plant-pathogen interactions (Xue *et al.,* 2022; Liang *et al.,* 2022 Li *et al.,* 2022). The nsLTP gene showed up-regulation during the elicitor-alone treatment in both cultivars, which showed up to one fold upregulation compared to control, *Alternaria solani* alone, and LaM pretreatment followed by *Alternaria solani*. The susceptible cultivar showed the minimum fold change in the *Alternaria solani* alone, LaM pretreatment alone, and LaM pretreatment followed by *Alternaria solani* compared to the control (**Fig. 12**.).

**Fig. 12.**
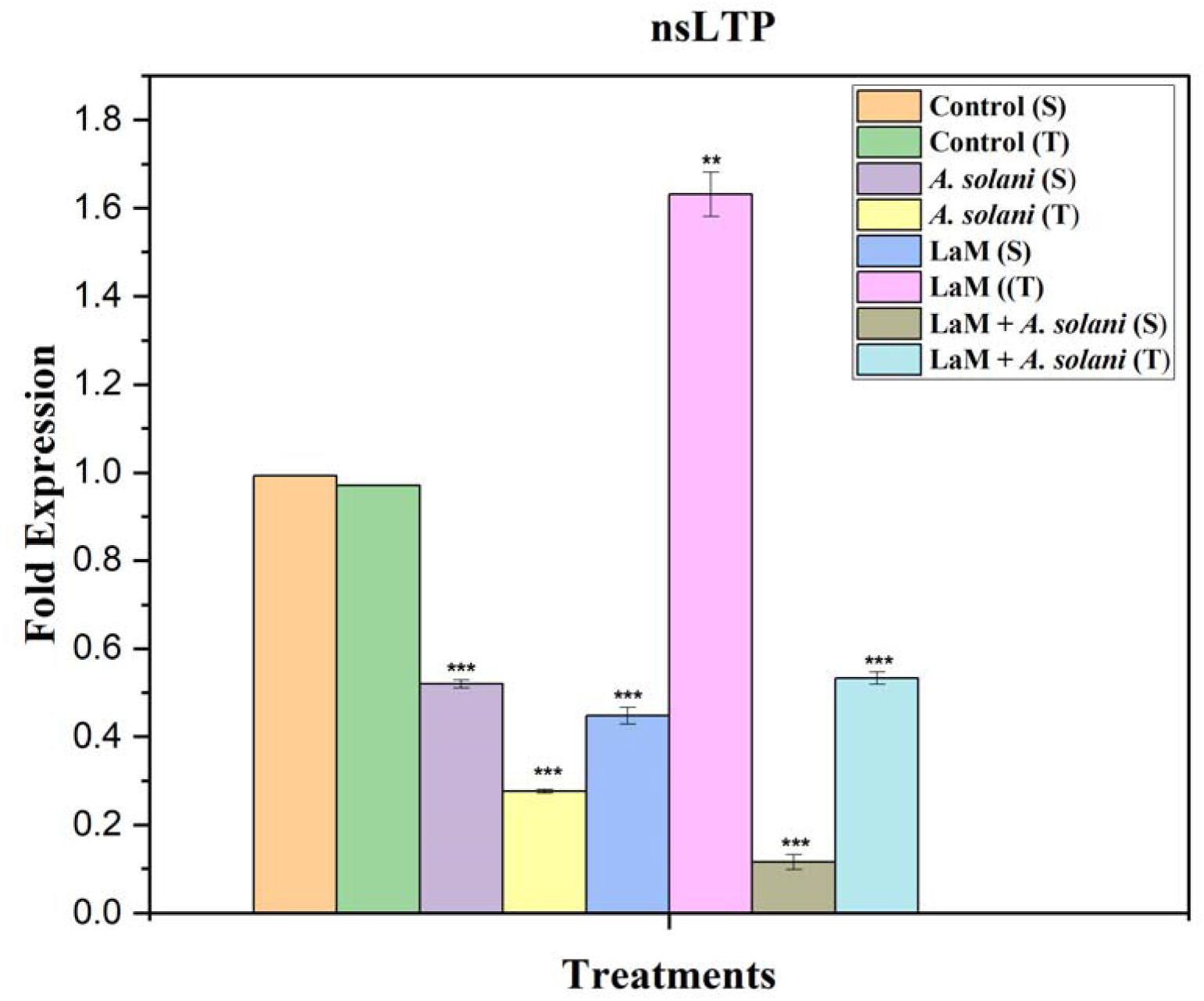
Relative expression profile of nsLTP gene in Control (S), Control (T), *A. solani* (S), *A. solani* (T), LaM (S), LaM (T), LaM followed by *A. solani* (S), LaM followed by *A. solani* (T) The relative expression ratio of each gene was calculated relative to its expression in the control sample. EF1α was used as an internal control to normalize the data. Error bars representing standard deviation were calculated based on three technical replicates for three independent biological replicates. Symbols; *, **, and *** indicate significant differences between control and treated samples (i.e. p< 0.05; p< 0.01 and p< 0.001, respectively) calculated using Student’s t test in OriginPro software.

SKP1 gene expression plays a vital role during biotic stress responses in plants. Studies have shown that SKP1-like proteins, such as ASK13 and PSK1, are up-regulated in response to abiotic stress (Rao *et al*., 2018; Hao *et al*., 2017). Similarly, the SKP1 protein is involved in the SCF complex, which is essential for protein degradation and plays a role in stress signaling pathways (Beji *et al*., 2019). Furthermore, SKP1 in *Dictyostelium discoideum* is involved in a hydroxyproline-linked pentasaccharide pathway, indicating its significance in stress responses (Choe *et al*.,2013). These findings collectively highlight the importance of SKP1 gene expression in mediating plant responses to biotic stress, suggesting its involvement in diverse cellular processes beyond protein degradation. The SKP1 gene showed up-regulation during the *Alternaria solani* infection in susceptible cultivars, which showed up to nine-fold upregulation in the *Alternaria solani* alone infected tomato plants compared to control and LaM alone pretreated tomato plants in the tolerant cultivar. In contrast, the maximum fold was observed in the tolerant cultivar in the control and other treatments (**Fig. 13**.).

**Fig. 13.**
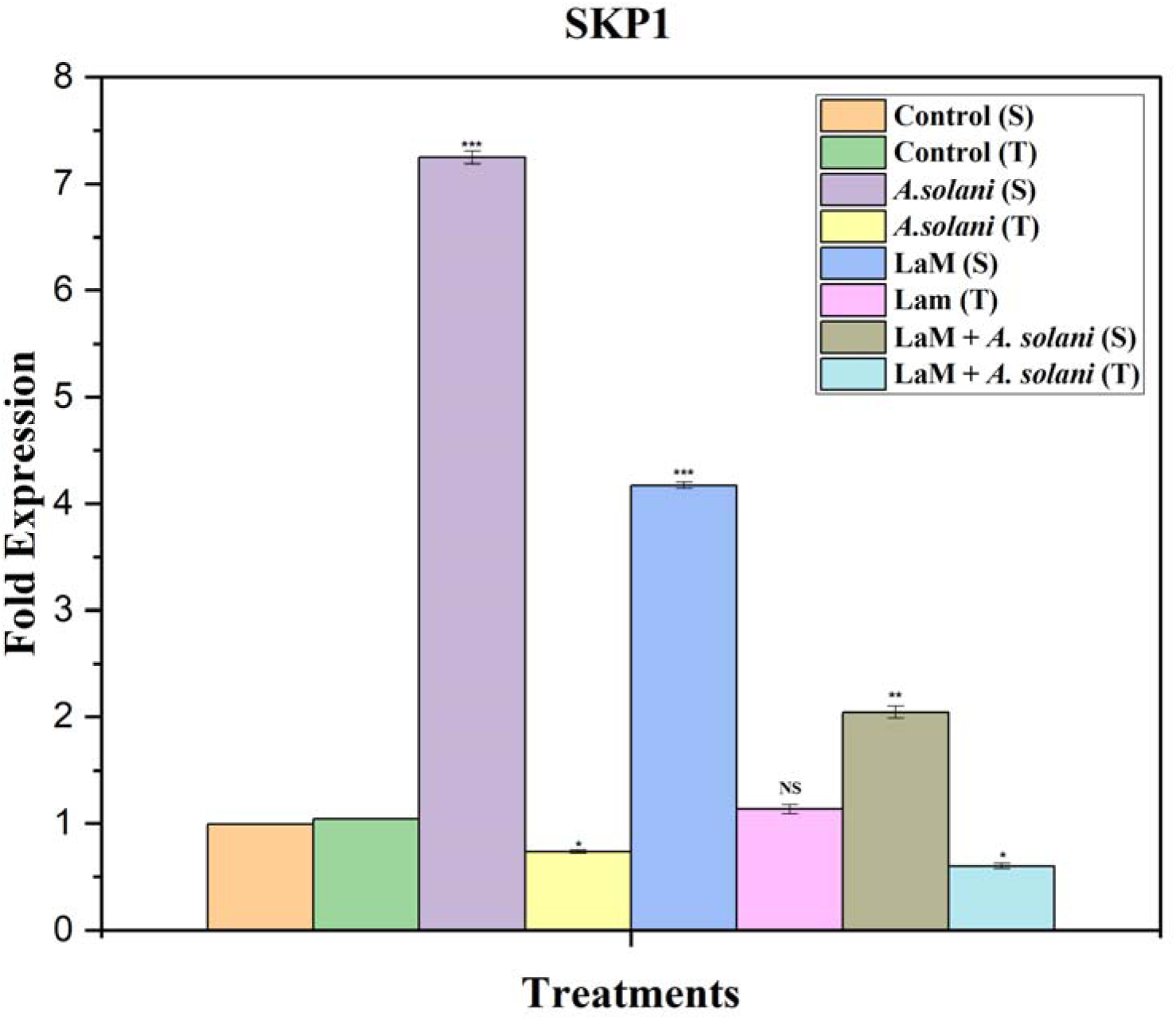
Relative expression profile of SKP1 gene in Control (S), Control (T), *A. solani* (S), *A. solani* (T), LaM (S), LaM (T), LaM followed by *A. solani* (S), LaM followed by *A. solani* (T) The relative expression ratio of each gene was calculated relative to its expression in the control sample. EF1α was used as an internal control to normalize the data. Error bars representing standard deviation were calculated based on three technical replicates for three independent biological replicates. Symbols; *, **, and *** indicate significant differences between control and treated samples (i.e. p< 0.05; p< 0.01 and p< 0.001, respectively) calculated using Student’s t test in OriginPro software.

The V-ATPase protein is essential for establishing proton gradients, aiding in ion sequestration, and enhancing stress tolerance (Zhang *et al*., 2014; Seidel, 2022; Gao *et al*., 2011). V-ATPase activity is vital for cell survival under stress conditions like salinity, drought, and other environmental challenges (Russak, 2011). The gene expression of V-ATPase can be induced by various stressors, including NaCl, NaHCO_3_, PEG, CdCl_2_, and ABA treatment (Dietz *et al*., 2001). Additionally, overexpressing V-ATPase genes has improved tolerance to salt, drought, UV, heavy metals, and extreme temperatures. Understanding the regulation of V-ATPase gene expression during biotic stress is crucial for improving stress resistance mechanisms and crop yield (Li *et al*., 2022). The V-type proton ATPase gene showed up-regulation during the pathogen infection in susceptible cultivars, which showed up to ten-fold upregulation in the *Alternaria solani* alone infection compared to control and LaM alone pretreated tomato plants. In contrast, the maximum fold change in the tolerant cultivar was evident in the control and other treatments (**Fig. 14**.).

**Fig. 14.**
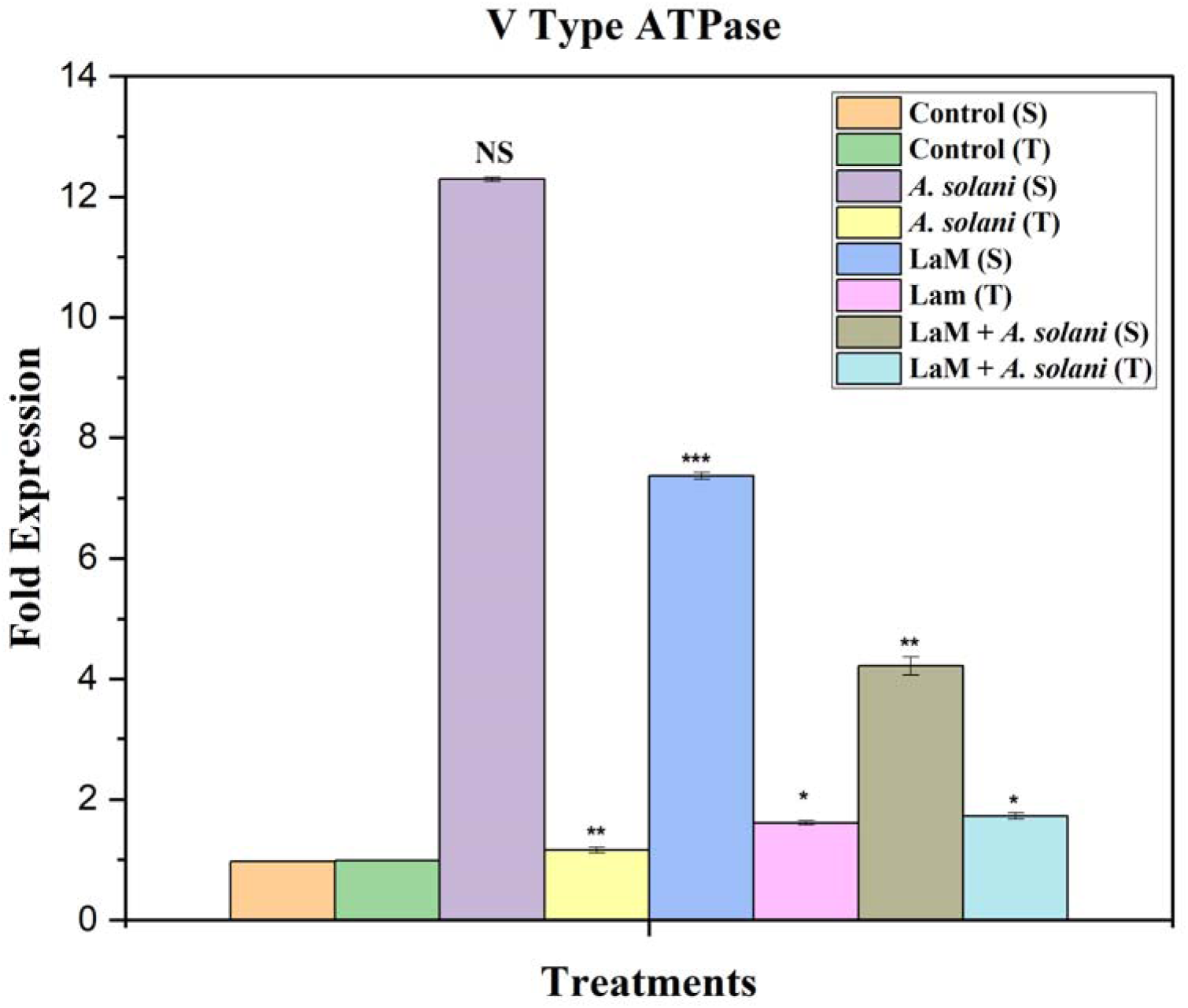
Relative expression profile of V type ATPase gene in Control (S), Control (T), *A. solani* (S), *A. solani* (T), LaM (S), LaM (T), LaM followed by *A. solani* (S), LaM followed by *A. solani* (T) The relative expression ratio of each gene was calculated relative to its expression in the control sample. EF1α was used as an internal control to normalize the data. Error bars representing standard deviation were calculated based on three technical replicates for three independent biological replicates. Symbols; *, **, and *** indicate significant differences between control and treated samples (i.e. p< 0.05; p< 0.01 and p< 0.001, respectively) calculated using Student’s t test in OriginPro software.

## CONCLUSION

In the present study, we have tested the efficiency of Laminarin in controlling the manifestation of *Alternaria solani* in *Solanum lycopersicum*. We elucidated that the elicitor-treated tomato plants followed by *Alternaria solani* showed significantly fewer symptoms in susceptible tomato cultivars when compared with *Alternaria solani* alone, infected susceptible and tolerant tomato cultivars. In order to study gene expression, transcriptome analysis was conducted for control, elicitor-pretreated, and elicitor-pretreated and/or pathogen-infected resistant and susceptible tomato cultivars. Differentially expressed genes of all the experimental leaves were analyzed, and it concluded that nsLTP, Aquaporin, Cytochrome P450, bZIP, and Heat shock protein 70 were upregulated and downregulated in different treatments. A few genes were selected from transcriptome analysis, and qRT-PCR validation was performed on all treatment conditions. Overall, the present investigation reveals that sulfated polysaccharide laminarin has the potential to induce defense-related genes, thereby offering tolerance against early blight disease caused by a fungal pathogen, *Alternaria solani.* From these results, it is inferred that pretreatment with Laminarin potentially reduced cell death in tomato plants caused by the *Alternaria solani* pathogen.

## ACKNOWLEDGEMENTS

The authors wish to thank The Director, Centre for Advanced Studies in Botany, University of Madras for providing the laboratory facility to carry out the experiments. The corresponding author also thanks the Tamil Nadu State Council for Higher Education (TANSCHE), Chennai, Tamil Nadu in India for providing financial support (RGP/2019-20/MU/HECP-0063-01).

